# Uncertainty in cardiovascular digital twins despite non-normal errors in 4D flow MRI: identifying reliable biomarkers such as ventricular relaxation rate

**DOI:** 10.1101/2024.09.05.611398

**Authors:** Kajsa Tunedal, Tino Ebbers, Gunnar Cedersund

## Abstract

Cardiovascular digital twins and mechanistic models can be used to obtain new biomarkers from patient-specific hemodynamic data. However, such model-derived biomarkers are only clinically relevant if the variation between timepoints/patients is smaller than the uncertainty of the biomarkers. Unfortunately, this uncertainty is challenging to calculate, as the uncertainty of the underlying hemodynamic data is largely unknown and has several sources that are not additive or normally distributed. This violates normality assumptions of current methods; implying that also biomarkers have an unknown uncertainty. To remedy these problems, we herein present a method, with attached code, for uncertainty calculation of model-derived biomarkers using non-normal data. First, we estimated all sources of uncertainty, both normal and non-normal, in hemodynamic data used to personalize an existing model; the errors in 4D flow MRI-derived stroke volumes were 5-20% and the blood pressure errors were 0±8 mmHg. Second, we estimated the resulting model-derived biomarker uncertainty for 100 simulated datasets, sampled from the data distributions, by: 1) combining data uncertainties 2) parameter estimation, 3) profile-likelihood. The true biomarker values were found within a 95% confidence interval in 98% (median) of the cases. This shows both that our estimated data uncertainty is reasonable, and that we can use profile-likelihood despite the non-normality. Finally, we demonstrated that e.g. ventricular relaxation rate has a smaller uncertainty (∼10%) than the variation across a clinical cohort (∼40%), meaning that these biomarkers have clinical usefulness. Our results take us one step closer to the usage of model-derived biomarkers for cardiovascular patient characterization.

**Highlights:**

- Digital twin models provide physiological biomarkers using e.g. 4D-flow MRI data
- However, the data has several non-normal uncertainty components
- For this reason, we do not know which biomarkers are reliable and clinically useful
- New method for data uncertainty and for calculation of biomarker uncertainty
- We identified several reliable biomarkers: e.g. ventricular relaxation rate

**Graphical abstract:** 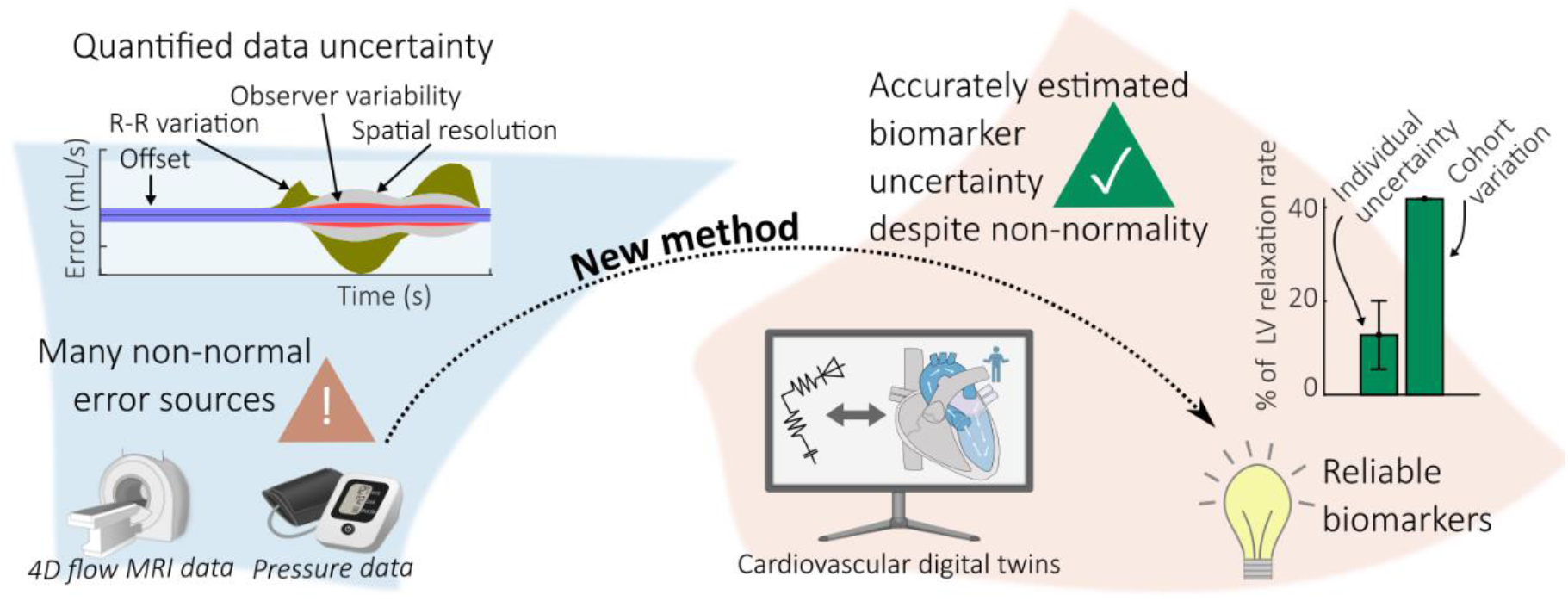

## 1. Introduction

Cardiovascular digital twins are made up of patient-specific mechanistic models for the cardiovascular system. Such twins have the potential to analyze patient-specific data to produce new patient-specific, model-derived biomarkers. For example, lumped parameter models have been used to analyze hemodynamic data, to predict biomarkers describing subject-specific hemodynamic parameters. Such biomarkers include e.g. left ventricular relaxation rate, aortic compliance, and left atrial pressure (Figure 1A). Studies predicting biomarkers have already been done in different conditions, including e.g. normal physiological conditions (Casas *et al*., 2017), stress (Casas *et al*., 2018), and disease (Khodaei *et al*., 2021; Harrod *et al*., 2021; Tunedal *et al*., 2023). Such biomarkers are highly valuable, but only if they are correct and if they have a low uncertainty (Hose *et al*., 2019; Viceconti *et al*., 2021; Musuamba *et al*., 2021).

**Figure 1.**
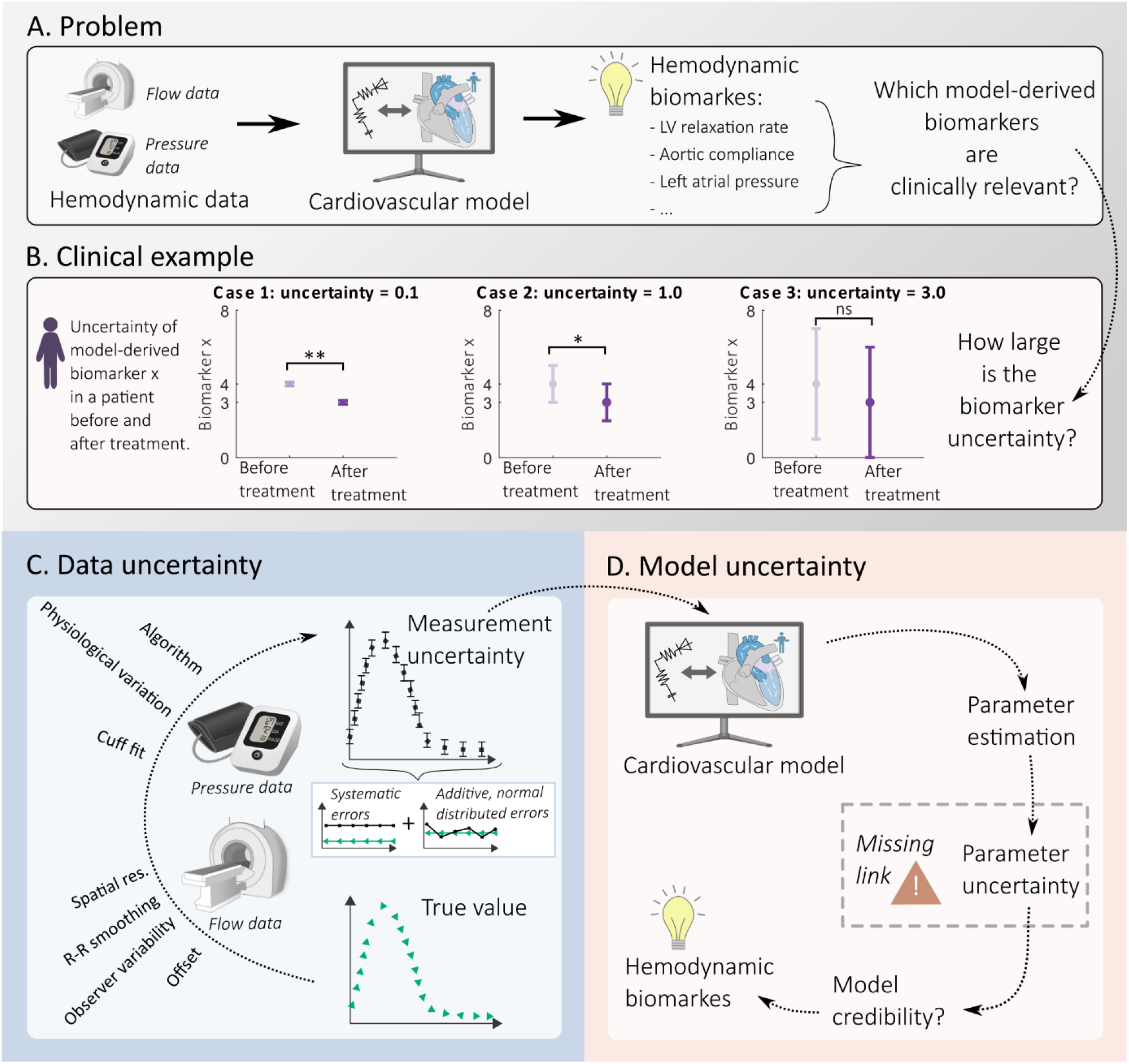
The uncertainty of the measurements used as training and validation data for a cardiovascular model propagates to the model-derived biomarker uncertainty. A) Hemodynamic biomarkers can be obtained from personalized cardiovascular models using hemodynamic data, but to know which of these biomarkers that are clinically relevant, one needs to know their uncertainty. B) If the biomarker uncertainty for a patient is small (Cases 1 and 2), one can see a significant change (*) in the biomarker x after treatment. However, if the uncertainty is too large (Case 3), the biomarker cannot be used and is thus not clinically relevant. Quantifying the data uncertainty (C) including all sources of errors for each measurement enables quantifying the model parameter uncertainty (D), which in turn enables evaluation of the credibility and usefulness of the parameters as hemodynamic biomarkers. Currently, the crucial parameter uncertainty estimation step is often missing, and the traditional estimation methods cannot handle non-normal systematic sources of errors in data such as offset or R-R smoothing.

The importance of this uncertainty is illustrated in Figure 1B. Consider a model-derived biomarker X, which has been measured in two conditions, e.g. before and after treatment. Before treatment the value was 4, and after 3, which may indicate a positive effect of the treatment. However, if that change is significant and meaningful wholly depends on the uncertainty of the biomarker: if the uncertainty is 0.1, the change is clear and significant, if the uncertainty is 1, it is much less clear, and if the uncertainty is 3, the observed change is of no significance. The same thing would apply if the biomarker was used to see differences between patients. This simple example illustrates that a biomarker is only useful if its uncertainty is smaller than the variations seen across time or a population. For many model-derived biomarkers, as in (Casas *et al*., 2017, 2018; Tunedal *et al*., 2023) (Figure 1A), we do not have a good estimate of the biomarkers’ uncertainties. This is because the underlying data uncertainty is largely unknown and has non-normal sources (Figure 1C) (Cedersund, 2012), which makes computation of the biomarker uncertainty challenging.

The uncertainty in hemodynamic data comes from a variety of different sources, and the sources depend on the type of data. For instance, blood flow data obtained from non-invasive 4D flow magnetic resonance imaging (4D-flow MRI) has several uncertainty sources, including e.g. background phase offset, observer variability, smoothing over several R-R intervals, and spatial resolution (Dyverfeldt *et al*., 2015) (Figure 1C). Many of these sources are systematic, i.e. they are not additive, normally distributed noise, but instead e.g. shift the entire curve up or down (Figure 1C). The combined impact, and the relative contributions, of all these error sources are not known. The same problem holds for other common hemodynamic data sources, including e.g. blood pressure data, which has its own sources of errors: e.g. cuff fit, physiological variations, and device-specific algorithms (Lewis, 2019; Stergiou *et al*., 2021; Liu *et al*., 2022). Also for blood pressure data, the combined impact and the relative contributions of the different error sources are not known. Therefore, we need to quantify the relative contribution of each error source to data, and propagate this uncertainty to a corresponding uncertainty in model-derived biomarkers.

The uncertainty of model-derived biomarkers from cardiovascular models has been estimated using various methods, each with their own critical limitations. Many studies use methods utilizing local or global sensitivity analysis (SA) as a basis for uncertainty quantification. These methods include e.g. Sobol sensitivity indices, stochastic sampling, or adaptive polynomial chaos expansion (Gul & Bernhard, 2015; Eck *et al*., 2016; Quicken *et al*., 2016; Pathmanathan *et al*., 2019; Randall *et al*., 2021). However, SA has problems accounting for nonlinear dependencies between parameters, which is common in cases of unidentifiability (Cedersund, 2012). These limitations are overcome by the profile likelihood (PL) methods, which works also in cases of unidentifiability (Cedersund, 2012; Kreutz *et al*., 2013; Lövfors *et al*., 2021). However, the PL method, just as most other methods, is based on a critical assumption that currently is not fulfilled in hemodynamic data: that the combined impact of all error sources on the data is known and can be described as a single error source that is additive and normally distributed. Because of methodological short-comings in dealing with non-normal error sources, in combination with a lack of knowledge regarding error sources in hemodynamic data (Figure 1C), the true uncertainty and consequently the clinical usefulness of hemodynamic model-derived biomarkers is unknown (Figure 1A).

In this paper, we introduce a new method to identify the uncertainty of model-derived biomarkers, to be able to identify which biomarkers that have a small enough uncertainty to be clinically useful. Therefore, we first quantify the relative contribution of all major error sources to 4D flow and blood pressure cuff data. Thereafter, we introduce a three-step method that comprises: Step 1) estimation of the combined data uncertainty including non-normal errors Step 2) estimation of biomarkers by fitting to data, Step 3) estimation of biomarker uncertainty with profile likelihood. To validate if our method works despite the non-normality of the error sources, we use simulated data from the resulting distributions of the combined data uncertainty. This allows us to compare the uncertainties obtained by our method with corresponding true biomarker values. Finally, we apply our new method to a clinical cohort, to compare the variation and uncertainty of six selected model-derived biomarkers, which allows us to identify some of the biomarkers that are well-determined enough to have clinical potential. Scripts to implement our method are provided at GitHub: https://github.com/kajtu/Uncertainty-estimation (Tunedal, 2024).

## 2. Method

The study was divided into three parts. First (Section 2.1), the data uncertainty was estimated by defining the most important sources of errors in 4D flow MRI, bSSFP MRI, and oscillatory blood pressure measurements. For each source of error, the magnitude and normality of the error were approximated based on previous studies. Second (Section 2.2), we introduce a new method to calculate the biomarker uncertainties. We evaluate this method using sampled data within the data uncertainty distribution from Section 2.1, to see if the new method can find the true confidence intervals for these biomarkers, despite the non-normal data uncertainty. The result validates our new methodology, consisting of the identified data uncertainty distributions and our method. Finally (Section 2.3), this new method is used to identify biomarkers that are reliable enough to capture variations across a clinical cohort.

### 2.1 Estimation of data uncertainty

The model training data for the investigated model (Tunedal *et al*., 2023) includes 4D flow MRI, 3D cine MRI short axis (SA) images, and brachial blood pressure. These data types amount to 12 different measurement variables, which in this text are denoted *y*_1_ − *y*_12_ where *i*=(1,12). For each of these 12 variables, the measurement uncertainty was defined and estimated. These uncertainties are calculated as a combination of different error sources. These sources can be classified into either 1) *random errors* (denoted *r*_*i*_ (*t*) with standard deviation *σ*_*i*_ (*t*)), where a new value drawn from a distribution is added specifically to each datapoint every time it is measured, or 2) *systematic errors* (denoted *s*_*i*_ (*t*)), where the same value is added to the datapoints every time the signal is measured. An example of a random error is observer variability, i.e. differences between e.g. how a region of interest is laid by different individuals. This error is random, since each time point in a time-series is impacted differently. An example of a systematic error is background offset, which adds the same offset to all time points in a measured blood flow curve. Another type of systematic error is RR variability. This error comes from the fact that 4D-flow data is an average of several heartbeats, which all have different RR-intervals compared to the average RR-interval. In simulation exercises (Section 2.2.1, 2.2.3), we analyzed this error and found that it can be approximated by a time-varying bias that is added to the entire time series. For each of the 12 variables, we identify each such error source (Table 2) and create a combined error (Table 3).

**Table 1.**
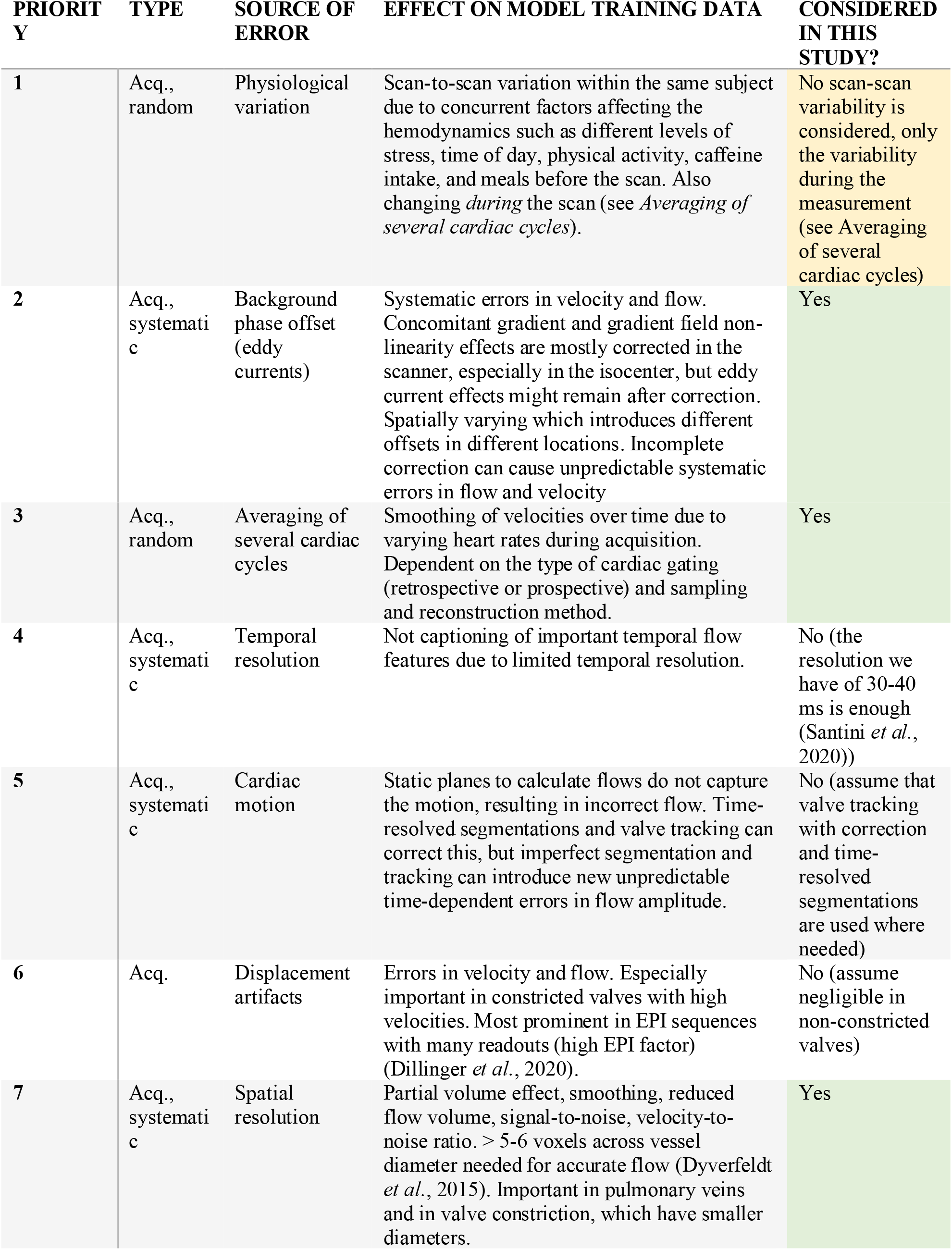
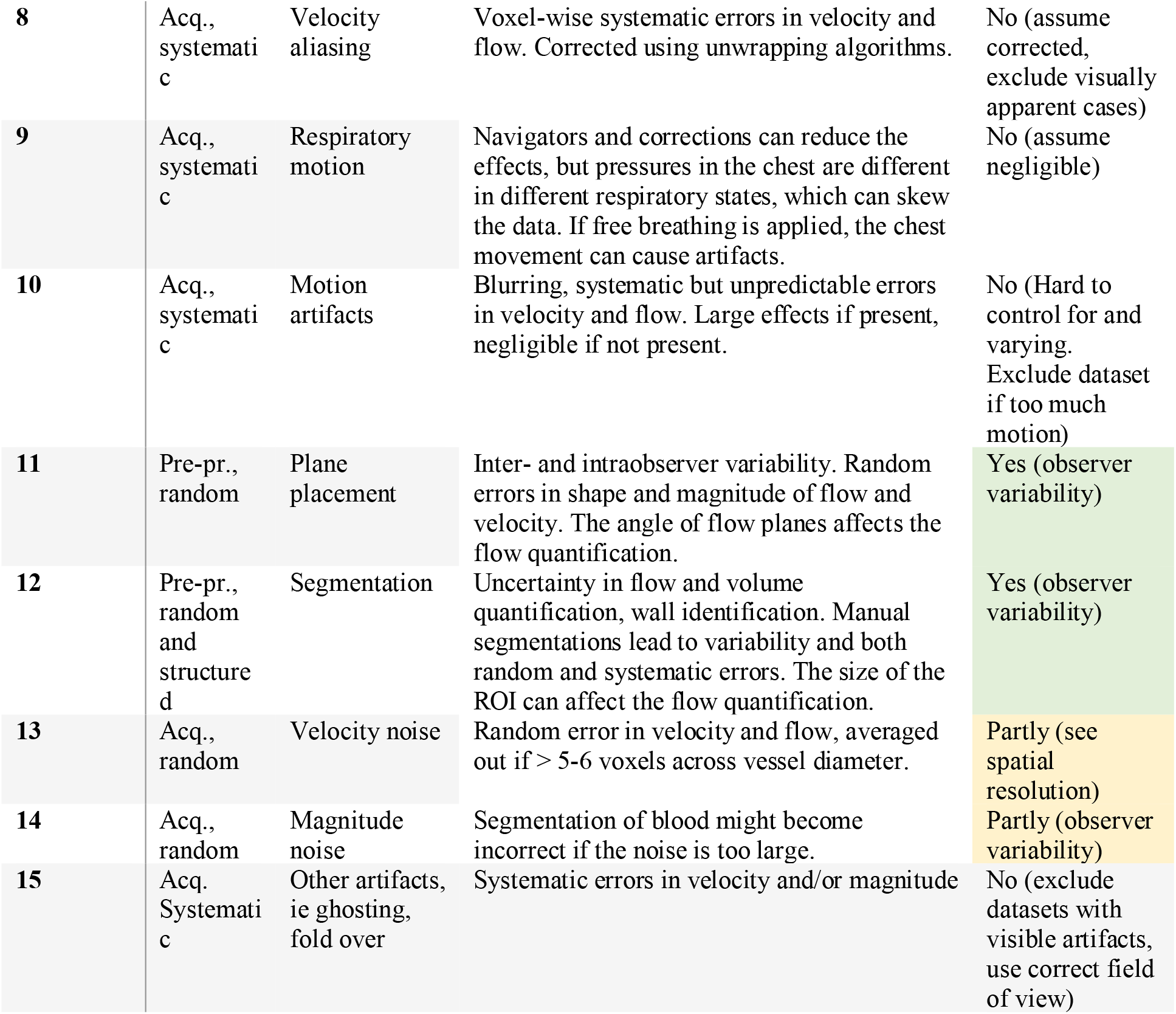
Sources of error in 4D flow MRI in prioritized order together with a description of their effect on the final data and if they are considered in this study or not. The priority is based on the estimated effect on the final data together with the probability of the error to occur. Type of error is defined as random or systematic and when in the imaging process it occurs: Acq. = acquisition, rec. = reconstruction, pre-pr. = preprocessing, an. = analysis. A more thorough description of each source of error is given in the text.

**Table 2.**
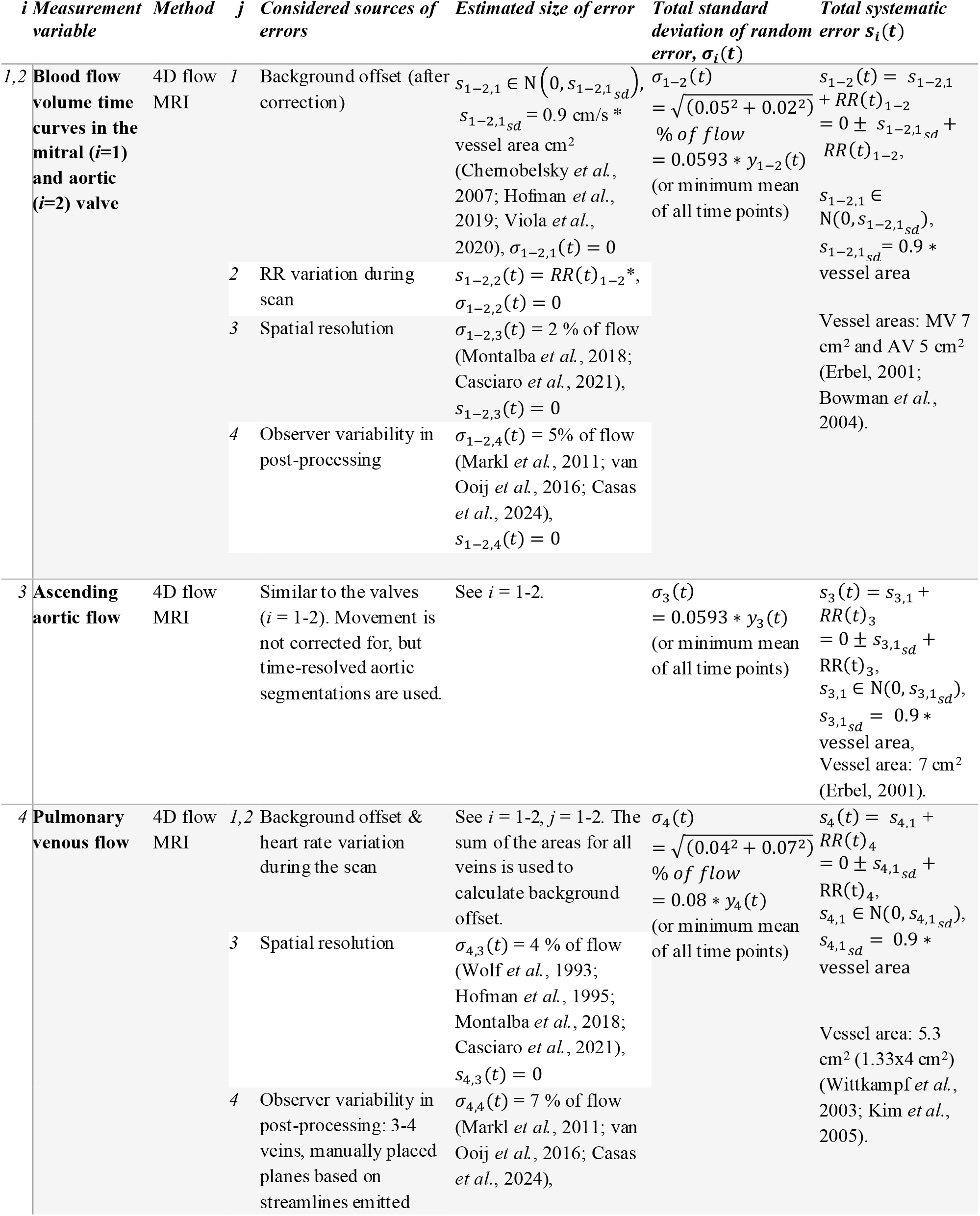
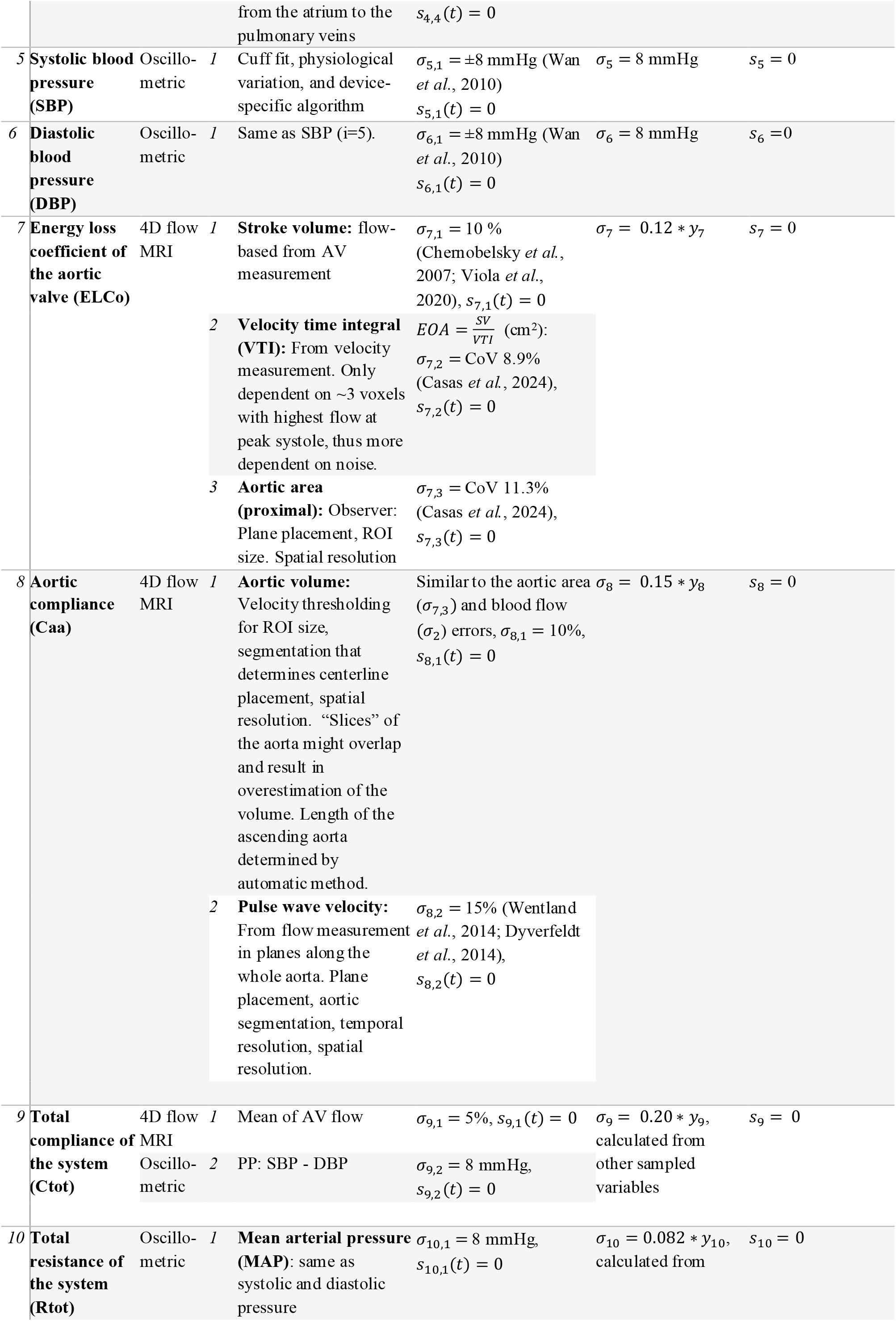
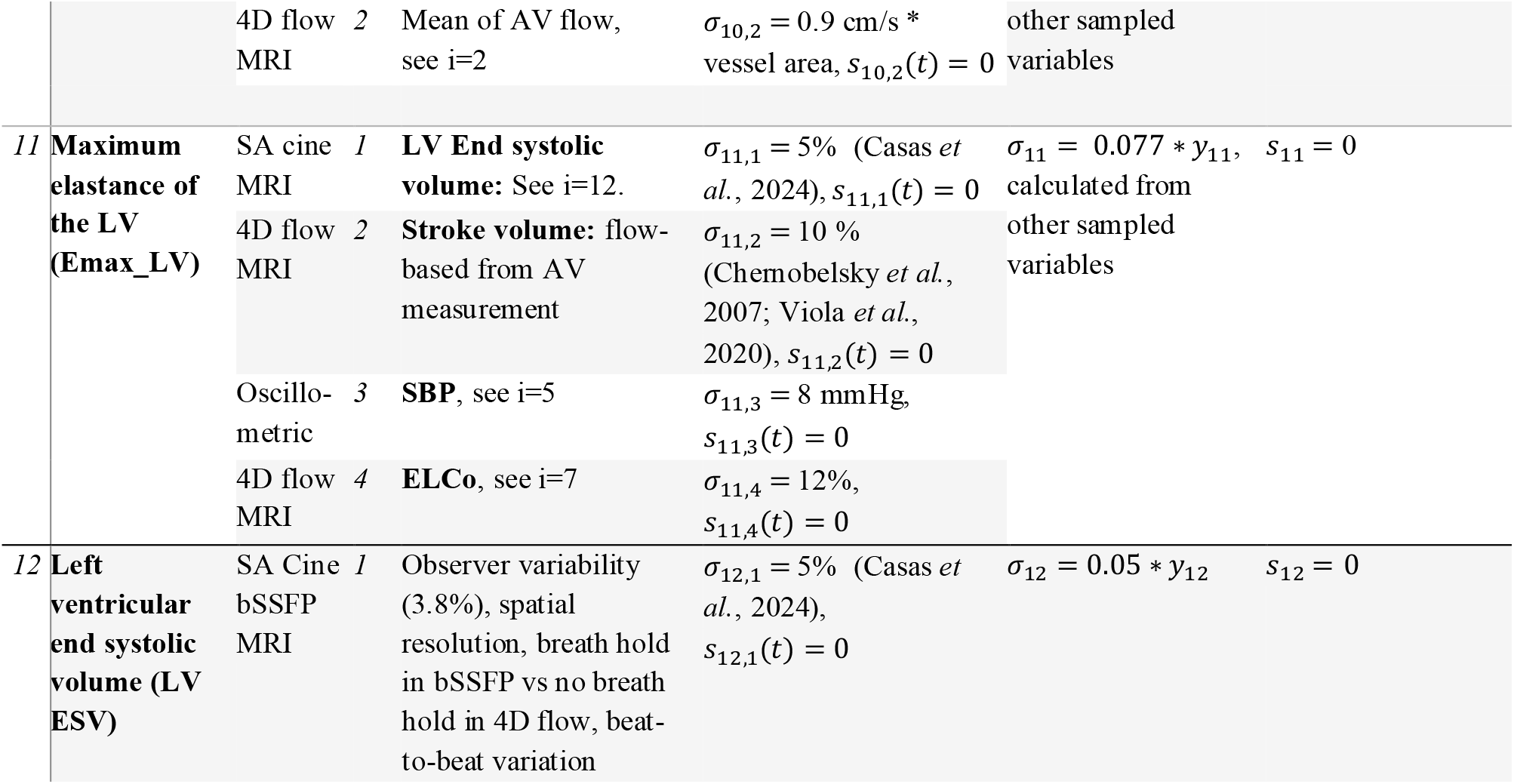
All 12 measurement variables i that are used to personalize the cardiovascular model, which method is used to measure them, which of the sources of errors j that are considered in this manuscript for each variable, and the size of each error source. The error sources are classified into systematic (s_ij_) and random (with standard deviation σ_ij_ (t)) (see Eq 5). For more details on each source of error, see Section 2.1. *RR(t) is the average of 400 random simulated heartbeats, see Section 2.1.1. MV: mitral valve, AV: aortic valve, AA: ascending aorta, PV: pulmonary veins, LV: left ventricle, RR: R-R interval, ie time of one cardiac cycle, sd: standard deviation.

**Table 3.**
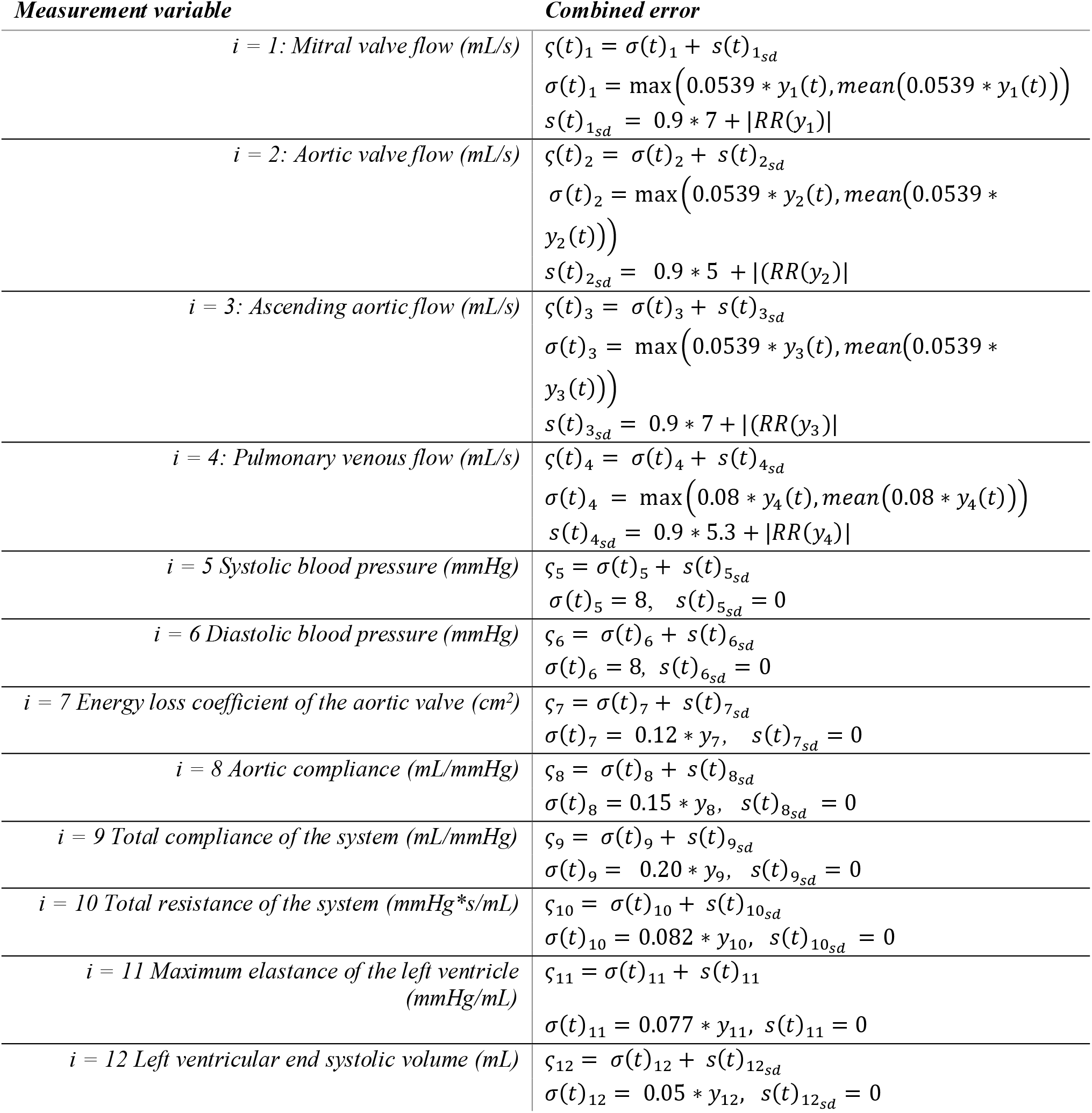
The combined error used in the parameter estimation and profile likelihood method, for each of the 12 measurement variables.

#### 2.1.1 Uncertainties in 4D flow MRI

Many of the 12 measurement variables stem from 4D flow MRI data (variables 1-4, 7-8, and 9-11). According to the 4D flow cardiovascular MRI consensus statement from 2015 (Dyverfeldt *et al*., 2015), the main sources of systematic errors in 4D flow MRI are background phase offset (eddy currents, concomitant gradient fields, gradient non-linearity) (Markl *et al*., 2012) and velocity aliasing. In addition to this, several other measurement features can potentially affect the accuracy of the measurements, such as spatial and temporal resolution, averaging of velocities within each voxel, averaging of the several cardiac cycles during the acquisition, and the noise level (Dyverfeldt *et al*., 2015). Finally, inter- and intra-observer variability can affect the results of the post-processing. The herein evaluated sources of error in 4D flow MRI are listed in Table 1, in a prioritized order, based on their estimated effect on the blood flow volume in the pulmonary veins, mitral valve, aortic valve, and aorta.

When estimating the size of the errors in 4D flow MRI, we only consider four errors: background offset, R-R variability, spatial resolution, and observer variability. We assume that all other errors and artifacts not considered here (such as motion, ghosting, and aliasing) are either negligible or corrected according to the 2023 4D flow consensus paper (Bissell *et al*., 2023). We also assume that datasets containing too large uncorrectable artifacts are excluded.

The magnitude of the first error, background offset, in the blood flow can be estimated by the conservation of mass principle within each dataset after correction. Viola *et al* found mass conservation differences of -8.96 to -0.25% with standard deviations of 5.9-11.8% for second-order polynomial background correction on 4D flow MRI data, which is around 8.6 mL with stroke volumes of 86 mL (Viola *et al*., 2020). After phantom correction, Chernobelsky *et al* found differences between the main pulmonary artery and the aorta of 7.1 ± 6.6 mL in 2D PC MRI data (Chernobelsky *et al*., 2007). These offsets in stroke volume around 8 mL correspond to flow offsets of 8 mL/s for 1s cardiac cycle length, and a velocity offset around 8/7 = 1.1 cm/s in a vessel with 3 cm in diameter. Hofman *et al* found remaining offsets of 0.1±0.5 cm/s compared to a phantom after interpolation-based offset correction (Hofman *et al*., 2019). Based on these three studies, we estimate the flow offset *after* correction as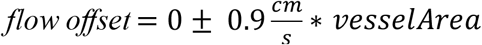. In this simplified estimation, we assume that correction is done and that the remaining offset is constant over the cardiac cycle.

The second error source, R-R variability, has previously been reported to be 0.095s or 10% during 4D flow MRI at rest before exercising (Macdonald *et al*., 2020), 1.3-7.2% during a breath hold for cardiac imaging (de Roquefeuil *et al*., 2013), and 61±22 ms i.e. 6.5% during 3D whole-heart coronary MR angiography (Roes *et al*., 2008). Here, we estimate the standard deviation of the R-R interval to 7%. To estimate the effect of R-R variation of blood flow curves, 400 heartbeats with 7% standard deviation and the true cardiac cycle length as mean value were drawn from a normal distribution, simulated with the cardiovascular model, and averaged (see Section 2.2.3). This is shown for one bootstrap in Figure 4 A-B iii where 10 of the 400 simulations, shown in lighter colors, are compared to the average in a dashed thicker line and to the true heartbeat in black. This is the process by which the error from RR variability is added to the simulated bootstrap analysis below (Section 2.2.3). When analyzing a real dataset, such a procedure is not possible, and one instead needs to consider the mean error from each error source. For RR variability, this mean error is obtained by the difference between the black and the dashed thicker line, shown in the green line in Figure 4 A-B, ii. As described in Section 2.2.1, we have created an empirical function to calculate this time-varying error for a given dataset. We classify the RR variability as a systematic error source, since it creates a time-varying systematic bias on the resulting blood flow curves (green line in Figure 4 A-B, ii).

The third error, from spatial resolution, is assumed to be relative to the magnitude of the flow. To simplify, we do not change the error with the resolution but assume that a reasonable resolution according to the consensus statement (Dyverfeldt *et al*., 2015; Bissell *et al*., 2023) is used. Thus, we estimate the effect of resolution as 2% of the flow in the mitral valve, aortic valve, and ascending aorta (Wolf *et al*., 1993; Hofman *et al*., 1995; Montalba *et al*., 2018; Casciaro *et al*., 2021). In the pulmonary veins with 3-5 voxels/diameter assuming a 3 mm isotropic resolution, we estimate the effect to be twice as large (Wolf *et al*., 1993; Pravdivtseva *et al*., 2022).

Finally, the fourth error, observer variability, has been reported between 2.8-8.9% for blood flow estimates (Markl *et al*., 2011; van Ooij *et al*., 2016; Casas *et al*., 2024) and is here estimated to be 5%.

#### 2.1.2 Uncertainties in bSSFP MRI

The sources of errors in balanced steady-state free precision (bSSFP) MRI are similar to the errors in 4D flow MRI. The error in end-systolic volume calculated from the short-axis cine of the left ventricle (measurement variable *i*=12) is estimated to be 5% based on a 3.8% segmentation observer variability and additional errors including spatial resolution, breath-hold in bSSFP sequences vs no breath-hold in 4D flow, and beat-to-beat variation (Casas *et al*., 2024).

#### 2.1.3 Uncertainties in brachial blood pressure measurement

Brachial blood pressure (measurement variables *i*=5-6) can be measured with several invasive and non-invasive techniques. Here, we focus on the non-invasive techniques of oscillatory blood pressure measurement since this is most practical in healthy controls. Blood pressure varies with the circadian rhythm and stress, is affected by movement and large heart rate variation, and often differs between measurements at home and the clinic (Stergiou *et al*., 2021). Additionally, oscillatory measurements depend on factors such as the device-specific algorithm, arm circumference, proper size of the cuff, and deflation rate (Lewis, 2019; Liu *et al*., 2022). In validation protocols, the mean absolute difference between test and reference measurements should be < 5±8 mmHg (ANSI/AAMI/ISO, 2018) although many validation studies report larger errors than 5 mmHg in practice (Wan *et al*., 2010). Here, we estimate the total error of automatic blood pressure measurement to be a random error with a standard deviation of 8 mmHg for both systolic (SBP) and diastolic (DBP) blood pressure.

#### 2.1.4 Uncertainties in calculated variables

The measurement variables Ctot, Rtot, and Emax_LV (*i=*9-11) are calculated based on other measurements (Eq 4), and the standard deviation and mean error for those three variables were therefore calculated via the uncertainty of the other measurements (see Section 2.2.2).

### 2.2 A new method to estimate uncertainty of model-derived biomarkers

The quantified data uncertainties were used to investigate the uncertainty of model-derived biomarkers. First, a new method for uncertainty estimation was introduced (Section 2.2.1), which handles the non-normality of several of the data sources. Second (Section 2.2.2), the cardiovascular digital twin model is introduced. Third, (Section 2.2.3), the method was validated using a bootstrap approach with simulated data based on the quantified uncertainty distributions from Table 2.

#### 2.2.1 A new method to estimate parameters with non-normal uncertainty

From the resulting 100 sampled datasets, we estimated the uncertainty of the biomarkers using a novel method, consisting of three steps (Figure 2):

**Figure 2.**
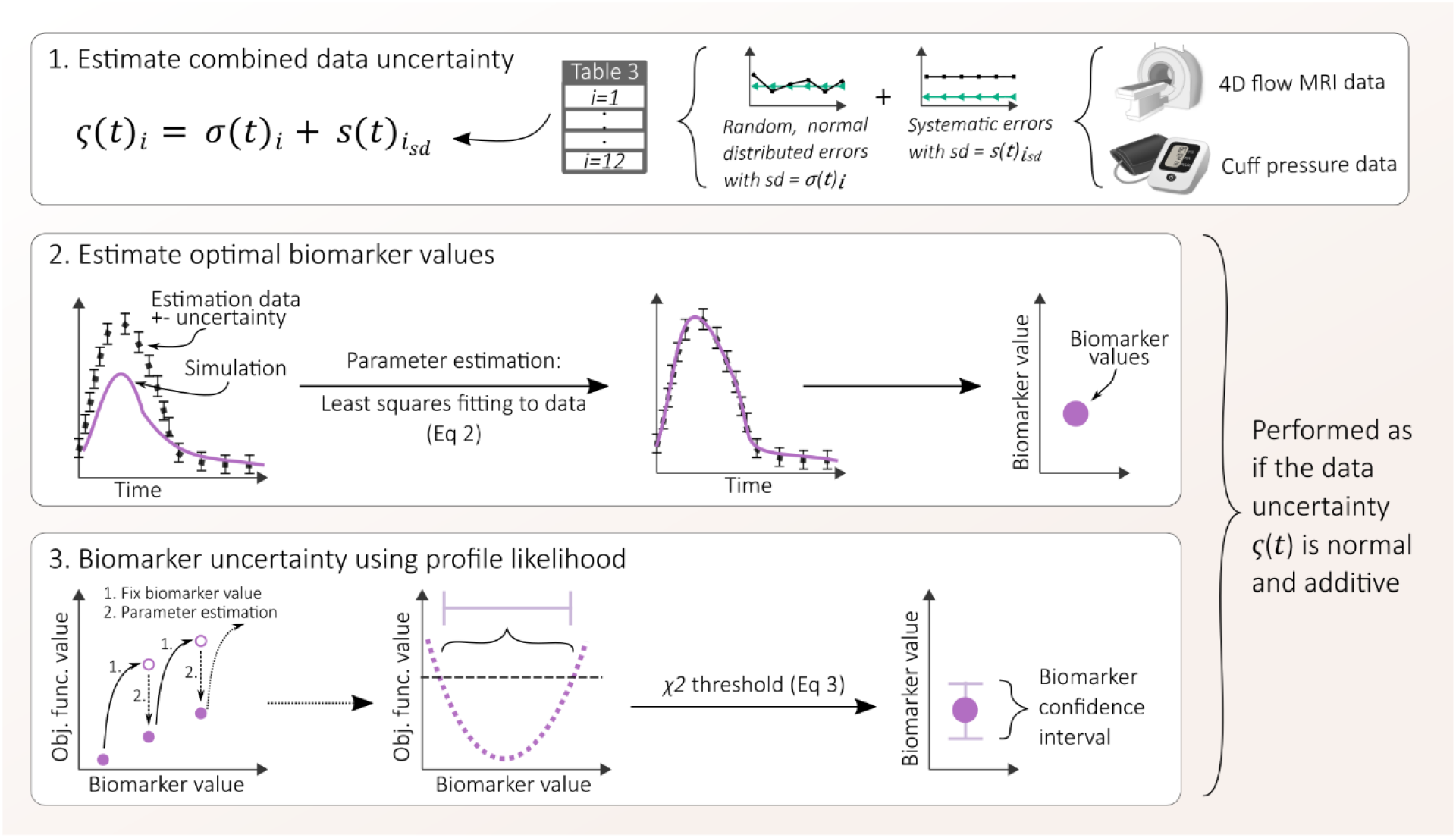
The method to estimate biomarker uncertainty using non-normal data. 1. All sources of errors in 4D flow MRI and cuff pressure data are estimated and classified as random and/or systematic. All standard deviations from all errors for each measurement variable i are then combined into an estimated data uncertainty (Table 3). 2. Using the estimated data uncertainty, the optimal biomarker values are estimated by fitting to data (Eq 2). 3. The biomarker uncertainty is then estimated using profile likelihood (Eq 3). For each biomarker, the biomarker value is fixed to one value at a time while all other biomarkers are estimated to minimize the difference between the simulation and the data. A 95% confidence interval is finally obtained using a χ^2^ threshold.

Step 1) Estimate the combined data uncertainty *ς*(*t*)_*i*_ using Table 3

Step 2) Estimate the optimal model-derived biomarkers by fitting to data using Eq 2

Step 3) Estimate the biomarker uncertainty with PL using Eq 3

The code to apply this method to your own data is available as Matlab scripts at https://github.com/kajtu/Uncertainty-estimation (Tunedal, 2024).

Note that steps 2-3 are done as if the data uncertainty was normal and additive. Let us now go through these steps in more detail.

##### Step 1: Estimate the combined data uncertainty

In this method, the uncertainty of the estimation data included both the distribution of systematic and random errors, resulting in a single combined error *ς*(*t*)_*i*_:

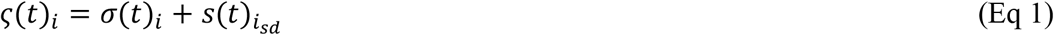

This final combined error uncertainty is described for each measurement variable in Table 3. Note that we add both the systematic and random errors together, instead of introducing the systematic errors as parameters. This reduces the computational cost by avoiding the estimation of additional model parameters and simulation of the RR error with several heartbeats for each single evaluation of the model. Note also that it is an estimate of the standard deviation of each error that is added to the combined error uncertainty in Eq 1, and not the full error *ε*_*i*_ (*t*) (Eq 5).

The handling of the RR variation error warrants some clarification. Since the smoothing due to RR variation in real, non-simulated data needs to be estimated without knowing the true flow curves, the size of the RR errors (i.e.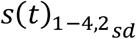) were estimated from each bootstrap using an empirical function *RR*(*y*_*i*_) that replicated the simulated RR errors obtained from the 100 bootstraps in Section 2.2.3 (available at https://github.com/kajtu/Uncertainty-estimation (Tunedal, 2024)). Note that *RR*(*y*_*i*_) is a function that can be both negative and positive, and that 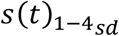 is positive, i.e. 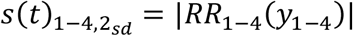.

For the calculated variables *i=*9-11, the combined uncertainty *ς*_9−11_ was estimated from all other bootstraps. For each bootstrap, the variable values were calculated, and the distribution of the difference between the 100 drawn calculated variables *y*_9−11_ and the true variables *y*^*τ*^ _9−11_ were used as an estimate of their uncertainty where the standard deviation is the random error *σ*_9−11_ and the systematic error was estimated to be 0.

##### Step 2: Estimate optimal biomarker values

In Step 2, the objective function of the weighted sum of squared residuals was minimized to estimate the biomarkers and provide a good starting point for the uncertainty estimations:

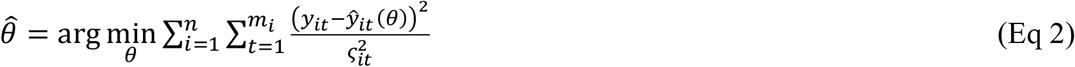

where *θ* is the parameter values for each dataset, 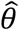 is the parameter vector theoretically maximizing the -2*log likelihood (−2LL), *y*_*it*_ is the measurement *i* at time point *t* out of *m*_*i*_ time points for measurement *i* and *n=12* measurements, and *ŷ*_*it*_(*θ*) is the corresponding model simulation. This is the classical least square estimation method.

In practice, the parameter estimation was done using several iterations of enhanced scatter search (ESS) in the MEIGO Matlab toolbox (Egea *et al*., 2014), starting with several runs of random guesses for the parameters and then using the previous best parameters as start guesses. The goodness of fit was evaluated with a χ^2^-test and a threshold value from the inverse cumulative density function 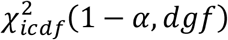 with the significance level *α* = 0.05 and degrees of freedom (dgf) equal to the number of measured datapoints. This test is done to make sure that the optimization has succeeded before going to Step 3.

##### Step 3: Estimate biomarker uncertainty with profile likelihood

In Step 3, the uncertainty was determined with profile likelihoods for all biomarkers (parameters) in the model, where the value of the parameter *p* is fixed while all other parameters in θ are re-estimated (Kreutz *et al*., 2013). The fixed value of *p* was initially set to a range with 50 steps based on the literature-based parameter bounds in (Casas *et al*., 2018; Tunedal *et al*., 2023): the lower bound divided by five to the upper bound times five. Starting at the optimum 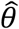, the value of *p* was first increased in steps until the upper bound times five was reached or until the -2LL reached above the value of 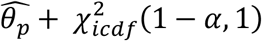, where *α* = 0.05. Then, the fixed value of *p* was decreased from the optimum until the boundary on the lower side was reached. At each step, the parameters were re-estimated in 7 repeated ESS. If needed, more optimizations were done at selected steps. Specifically during the method evaluating using bootstraps with known true values, smaller steps were taken close to the true value of *p* if needed to ensure that small enough steps was taken to accurately determine if the true value was within the model uncertainty. In general, smaller steps were taken closer to the defined limit of the - 2LL. Finally, the confidence interval for each parameter *p* was determined as all values of *p* that fulfills Eq 3:

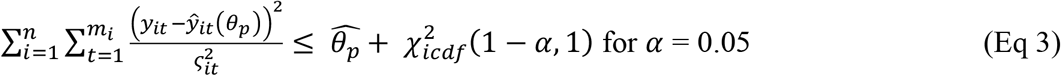

#### 2.2.2 The personalized cardiovascular model

The uncertainty estimation method was applied to a personalized cardiovascular model describing the left heart and ascending aorta (Figure 3A) (Casas *et al*., 2018; Tunedal *et al*., 2023). The model is a simple lumped parameter model and contains 28 parameters that are estimated, and 11 parameters that are kept constant. The parameters are estimated based on 4D flow MRI-derived blood flow volume curves in the pulmonary veins, mitral valve, aortic valve, and ascending aorta, systolic and diastolic blood pressure, and five data-based parameters. The data-based parameters are the aortic compliance Caa, the total compliance Ctot, the total resistance Rtot, the energy loss coefficient in the aortic valve ELCo, and the maximum elastance in the left ventricle Emax_LV, which are calculated as follows (Eq 4 a-e):

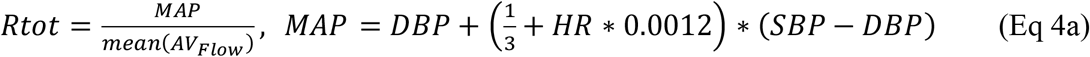

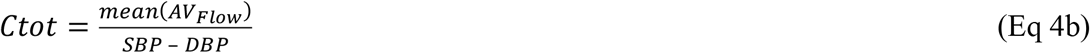

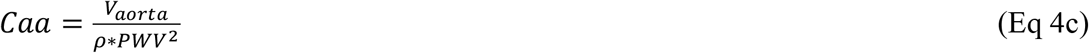

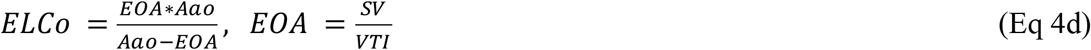

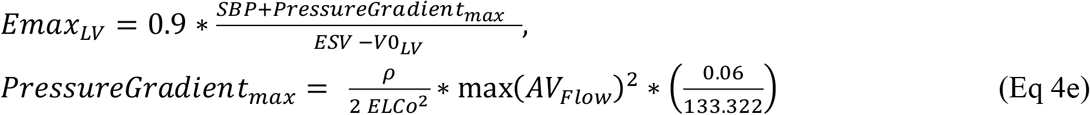

where *AV*_*Flow*_ is the measured blood flow through the aortic valve, SBP and DBP are the systolic and diastolic blood pressure, HR is the heart rate, *V*_*aorta*_ is the volume of the ascending aorta, *PWV* is the pulse wave velocity in the whole aorta, *ρ* is the density of the blood, *SV* is the stroke volume of the aortic valve, *VTI* is the velocity time integral in the aortic valve, *ESV* is the end-systolic left ventricular volume, and *V0*_*LV*_ is the unstressed left ventricular volume. *ρ* and *V0*_*LV*_ are constants in the model, set to 10 mL and 1.06 g/mL. A description of the 28 estimated parameters is available in the supplementary (Table S1).

**Figure 3.**
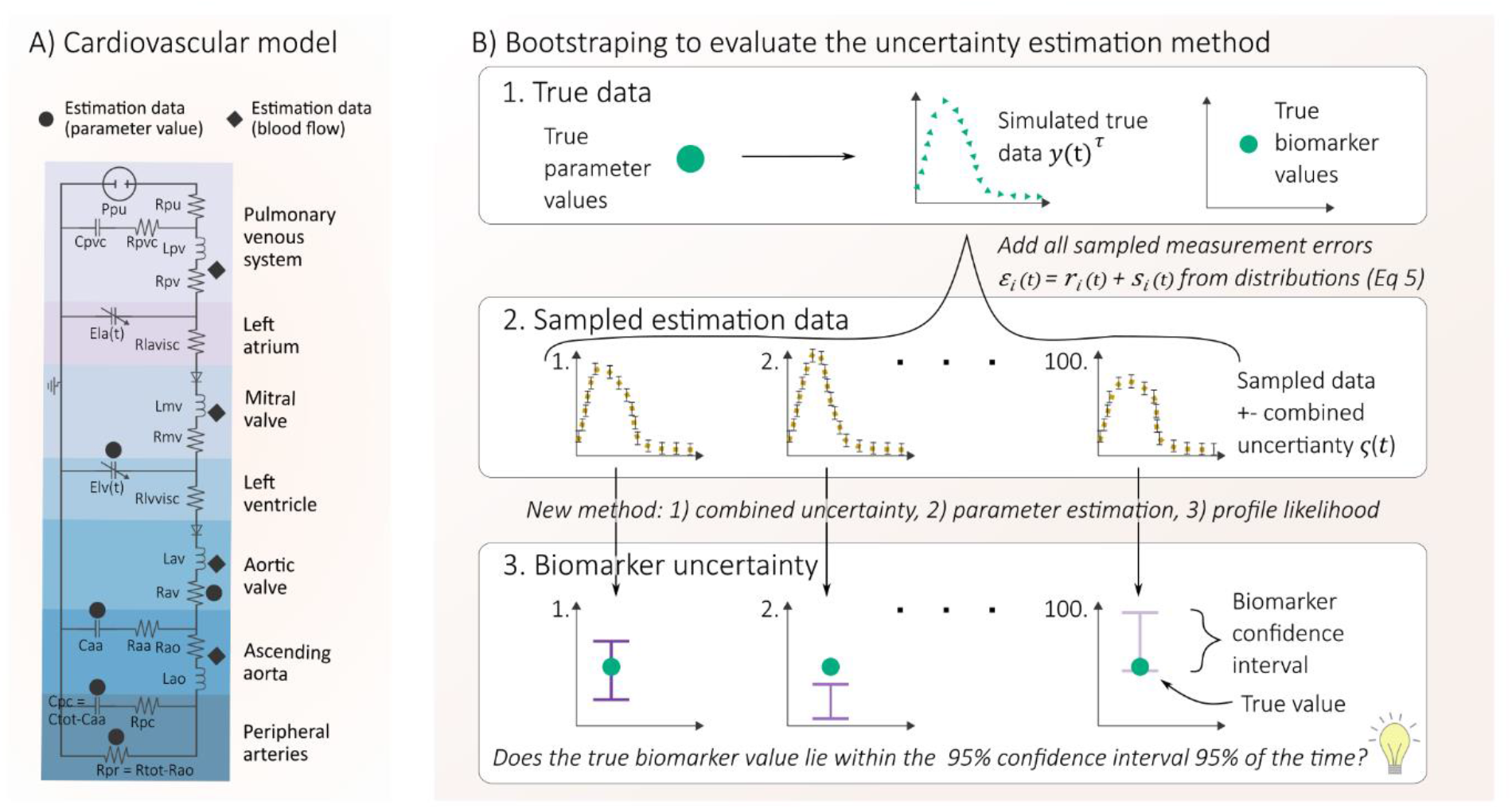
A) The cardiovascular model. B) Method to evaluate the novel method including profile likelihood-derived uncertainty of model-derived biomarkers. 1. A simulated true dataset was created with known true biomarker values. 2. 100 sampled datasets were generated by adding measurement errors from the quantified distribution of random (r_i_ (t)) and systematic errors (s_i_(t)) (Eq 5) and an estimated combined measurement uncertainty ς(t)_i_ (Table 3, Eq 1). 3. Each sampled dataset was used to estimate the uncertainty of model-derived biomarkers using our novel method. Finally, the biomarker confidence intervals were compared to the true biomarker values.

**Figure 4.**
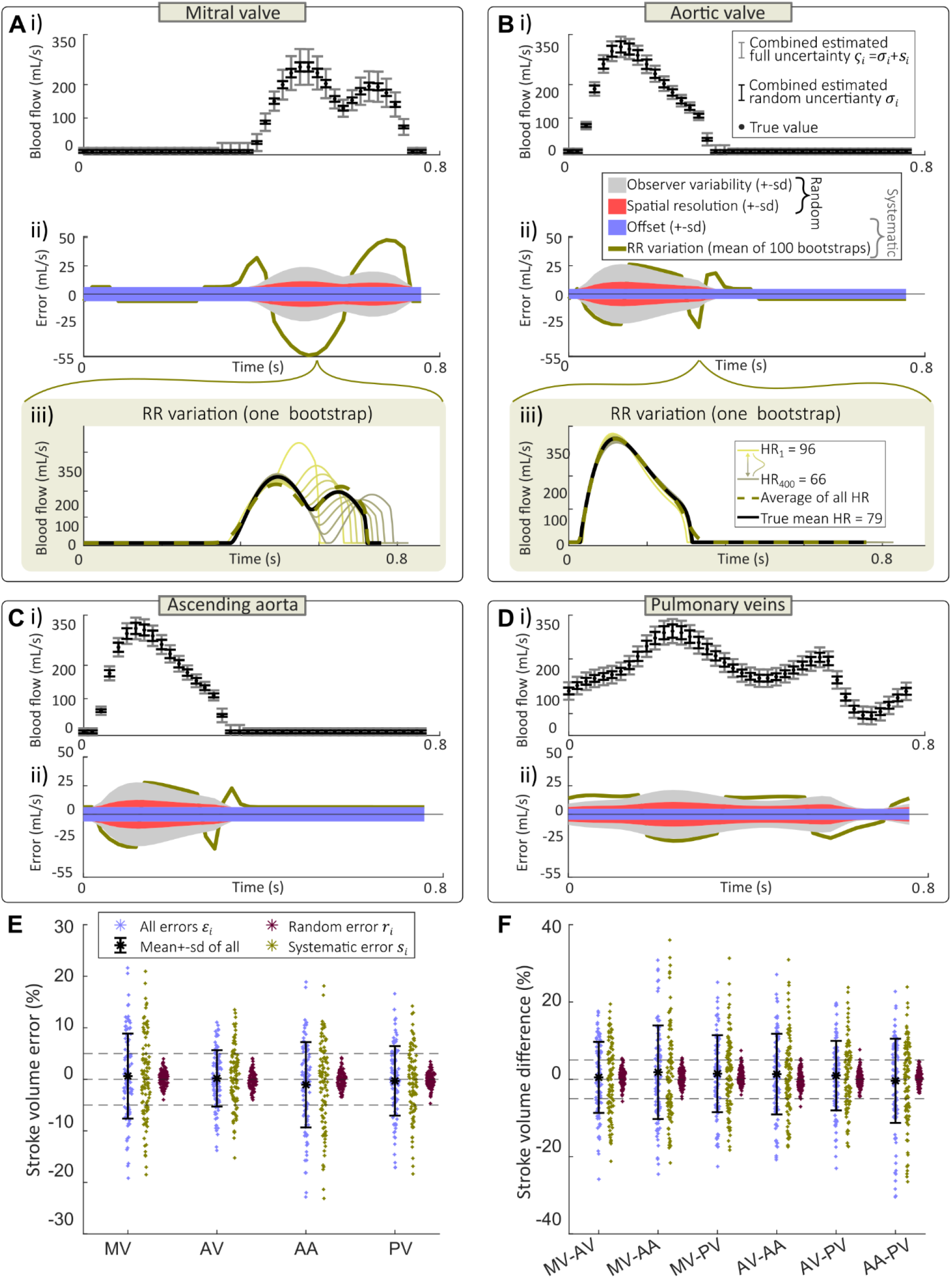
Estimated measurement uncertainty for 4D flow MRI. A-D) The size of different sources of errors in the blood flow curves used as training data for the cardiovascular model. Part i) shows the standard deviation of the estimated combined errors in relation to the mean values of the blood flow. The standard deviation σ_i_ of the combined random errors (spatial resolution and post-processing) are shown in black error bars. Grey error bars show the full estimated combined ς_i_ when including both random and systematic errors during parameter estimation for the model. Part ii) shows the size of the individual errors added on top of each other: the mean systematic smoothing due to R-R variation of all 100 bootstraps (green), the standard deviation (sd) of the systematic background offset (blue), sd of random observer variability (grey), and sd of random errors due to spatial resolution (red). Since the size of the errors due to offset, spatial resolution, and post-processing varies randomly, the standard deviation of each error is shown here. The RR smoothing is estimated by simulation of a distribution of 400 heart rates for each sample, and taking the average of the resulting blood flow curves, as shown for one bootstrap in A-B iii) where 10 examples of the 400 simulated heartbeats are shown in light lines, the average of the 400 simulations in the dashed green line, and the simulation of the true heart rate in the black solid line. The difference between the average and the true simulation is the RR error RR(t), and the mean RR error of all 100 bootstraps is shown in the green line in ii). E) The error in stroke volume in the mitral valve (MV), aortic valve (AV), ascending aorta (AA), and pulmonary veins (PV) when comparing the 100 sampled datasets to the true simulated stroke volume. F) Internal differences in stroke volume between different places in the imaging volume. In both E and F, the error when adding all measurement uncertainties is shown in blue to the left, the error when only adding systematic errors is shown in green in the middle, and the error when only adding random errors is shown to the right in red. The dashed lines represent a 5% error.

#### 2.2.3 Evaluation of the uncertainty estimates using bootstraps with known true values

To evaluate if our method (Section 2.2.1, Figure 2) could correctly estimate the uncertainty of the model-derived biomarkers despite the non-normal error sources in data, we used the estimated data uncertainty distributions in Table 2 to draw 100 bootstraps (Figure 3B 1-2). Finally, the new method was used to estimate 95% confidence intervals for all biomarkers using the 100 sampled estimation data, and it was evaluated how many times the true biomarkers lay within the confidence intervals (Figure 3B, 3).

The bootstraps are obtained by first creating one simulated true dataset (Figure 3B, 1). The simulated true dataset 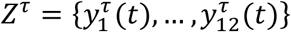, consisting of 12 model outputs corresponding to the 12 measurement variables, was created from a model simulation with the cardiovascular model, where the true model biomarkers were known. The model simulation was done with parameters estimated from MRI and blood pressure data in a healthy control from (Tunedal *et al*., 2023) to replicate a realistic physiology. Second, errors are added to the simulated “true” values *y*^*τ*^ _*i*_ (*t*) for each measurement *i* according to the distribution of the error sources identified in Table 2, creating 100 bootstraps (Figure 3B, 2). For each bootstrap *Z* = {*y*_1_ (*t*), …, *y*_12_ (*t*)}, 12 measurement variables *y*_*i*_ (*t*) were created by drawing errors from the uncertainty distributions *ε*_*i*_ (*t*) in Eq 5.

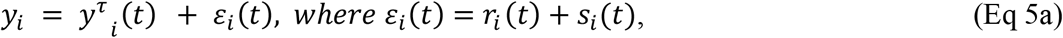

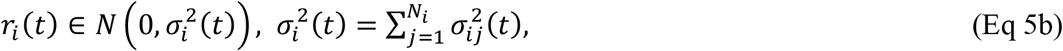

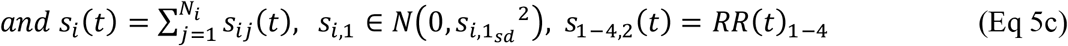

where each measurement variable *y*_*i*_ has *N*_*i*_ number of error sources, that are classified as random (*r*_*ij*_ (t)) and/or systematic (*s*_*ij*_ (*t*)), and that all are combined into the full measurement error *ε*_*i*_ (*t*). All error distributions are based on the estimated data uncertainty described in Table 2 and Section 2.1.

The random errors (Eq 5b) are combined by summing their variances 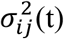, and the combined error *r*_*i*_ (t) is randomly sampled within its normal distribution with the standard deviation *σ*_*i*_ (t) in each time point *t*.

The systematic errors *s*_*ij*_ (*t*) are summed into *s*_*i*_ (t) (Eq 5c). Most systematic errors are offset errors that are the same for all time points and are drawn from a normal distribution 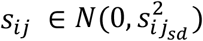. The only exception is the 4D flow MRI-derived flow curve variables 1-4 which have one time-dependent error source *s*_1−4,2_(*t*) = *RR*(*t*)_1−4_. RR(t) was sampled by simulating 400 heartbeats with cardiac cycle lengths T drawn from the distribution *N*(*T*_*true*_, 0.07 * *T*_*true*_). All resulting flow curves were averaged, and the error *RR*(*t*)_1−4_ was calculated as the difference between the true simulation, with cycle length *T*_*true*_, and the averaged value. When changing the heart rate in the model simulations, the model parameters k_diast_LV, onset_LA, and Emax_LV governing the ventricular and atrial contraction and relaxation were changed from the original value *pvalue*_*orig*_ with the formula *pvalue* = *T* * *k* + (*pvalue*_*orig*_ − *T* * *k*) and the k-values -0.55, 0.1, and -6, to achieve a change in systolic and diastolic time similar to (Gemignani *et al*., 2008) and to achieve the expected decrease in E/A ratio with increased heart rate (Yamamoto *et al*., 1993; Chung & Kovács, 2006).

### 2.3 Biomarker uncertainty in a clinical dataset

To see which model-derived biomarkers that are useful in practice, the method described in 2.2.1 was applied to a few selected biomarkers in a subset of subjects from a real clinical dataset from (Tunedal *et al*., 2023). Six biomarkers were selected for in-depth investigation of reliability: m2LV, Cpvc, and ksystLV that in Tunedal *et al* 2023 all were significantly different between any of the groups, Emax LA to include an atrial biomarker that was different between groups, Rao that had the strongest correlation with systolic blood pressure during MRI, and Caa that was the only biomarker that correlated with home blood pressure. The uncertainty of these biomarkers was estimated for 3 controls with a good fit to data and 3 subjects with hypertension and diabetes with a good fit to data. To remain within physiological bounds during PL, none of the parameters were allowed to be less than 0, Cpvc was kept <= 100, and ksystLV <= 200. Pairwise differences between subjects were considered significant and marked with * if there were no overlaps of their 95% confidence intervals. The resulting individual uncertainty in percent was evaluated for each subject and each biomarker as 100*(*sd*)/*p_mean*, where sd is the standard deviation of the individual uncertainty, and *p_mean* is the mean biomarker value corresponding to the best fit to data across all subjects. The cohort variation was calculated for each biomarker as 100*(*sd_all)*/*pbest_mean*, where *sd_all* is the sd of the biomarker value corresponding to the best fit to data across all subjects for that biomarker.

## 3. Results

First, all major error sources in 4D flow MRI (Section 3.1.1), cuff pressure (Section 3.1.2), and the calculated variables (Section 3.1.3) were quantified. Thereafter (Section 3.2), our new method (Section 2.2.1, Figure 2), was validated using a bootstrap approach (Section 2.2.3, Figure 3) using the quantified data uncertainties. Finally, the new method was applied to a clinical dataset to reveal reliable biomarkers with small uncertainty (Section 3.3).

### 3.1 Quantified data uncertainty

#### 3.1.1 Impact of different sources of errors in 4D flow MRI

For 4D flow MRI, we have performed an extensive literature analysis, combined with other reasoning, to identify 4 different error sources: background offset, R-R variation, spatial resolution, and observer variability (Table 2). These error sources yield two types of errors in data: a) 7-12 % random uncertainty of 4D flow MRI-derived blood flow due to observer variability in post-processing and spatial resolution, and b) systematic errors due to R-R smoothing and background offset.

We have visualized the estimated error contributions in Figure 4. This figure shows the combined error used in model parameter estimation (Figure 4 A-D, i), as well as the magnitude of the individual error distributions used to draw all of the bootstrapped estimation data, added upon each other to show their combined size, (Figure 4 A-D, ii), and the effect of these errors on the stroke volume (Figure 4 E-F).

A number of observations can be made. First, the size of the random errors (inner error bar in each time point) is larger than the systematic errors for most time points, especially in the aortic valve, ascending aorta, and pulmonary veins (Figure 4 B-D). However, since the errors are random, the combined effect of random errors on stroke volume is relatively small – less than 5% (Figure 4E).

Second, the effect of the systematic R-R smoothing is largest on the mitral flow (Figure 4B), where the E/A ratio is affected with a reduced early peak and an increased late peak and a mean effect of 0.5 mL/s for all time points compared to 0.1 and 0.2 mL/s for the aortic valve and ascending aorta. The smoothing effect is also visible in the pulmonary veins with a mean effect of 0.5 mL/s. The reason for this variation in the size of the RR smoothing can be seen in Figure 4 A-B iii, where and example of the averaging of 400 RR intervals is shown. When the heart rate increases, in this example up to 96 bpm, the length of diastole is shortened. This leads to the A-peak from the atrial contraction merging with the early E-peak from the ventricular relaxation in the mitral flow (Figure 4A iii). For lower heart rates, in this case down to 66 bpm, the opposite happens. This change in dynamics of the flow in the mitral valve is not completely linear, which results in the underestimation of the early peak and overestimation of the late peak when averaging over all 400 heartbeats for this specific heart rate interval. In the aortic valve and the ascending aorta, on the other hand, the RR effect is very small since the time of systole barely changes with the heart rate (Figure 4 B, C ii).

Third, the effect of background offset, which in the figure is set to ± one standard deviation, is in many time points very small, only a few mL, compared to the total blood flow (Figure 4 A-D ii). Due to the difference in vessel areas, the size of the offset error also varies between the four locations. However, the offset effect on the stroke volume is the largest of all sources of errors with up to 20% stroke volume error in the worst case of the 100 sampled datasets, and in general 0.43 ± 8.2 % error in mitral valve stroke volume for systematic errors only compared to 0.19 ± 1.51 % error for the random sources of error (Figure 4E, Table S2). Also in the stroke volume error between flows (Figure 4F), the background offset causes the main part of the error while the random error is smaller. The total stroke volume error between flows, i .e. error in mass conservation, is around -0.28-1.51 ± 8.9-11.9 % and up to 30% in some cases (Table S2).

#### 3.1.2 Impact of different sources of errors in brachial cuff pressure

For brachial cuff pressure, the errors due to cuff fit, physiological variation, and device-specific algorithm were estimated to 0±8 mmHg. Examples of 10 out of 100 resulting sampled measurements of systolic and diastolic pressure are shown in Figure 5 E.

**Figure 5.**
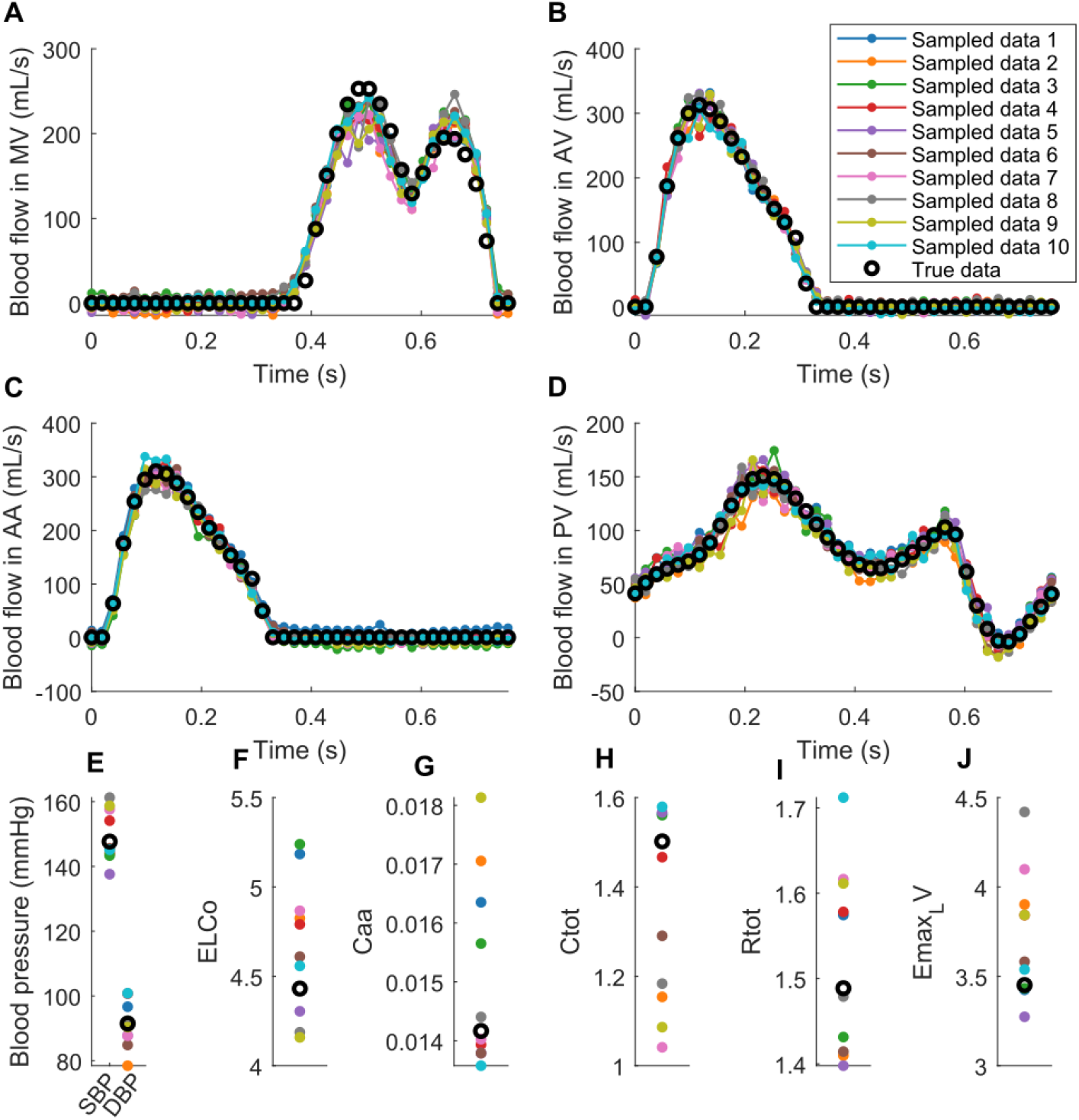
Examples of 10 of the 100 datasets sampled using bootstrapping based on a simulated true dataset (black circles) and the estimated size and distribution of the measurement uncertainty. Each dataset is represented by one color and contains measurements of blood flow in the A) mitral valve, B) aortic valve, C) ascending aorta, and D) pulmonary veins, as well as E) systolic and diastolic blood pressure, F) energy loss coefficient of the aortic valve, G) compliance of the ascending aorta, H) total compliance, I) total resistance, and J) maximum elastance of the left ventricle.

#### 3.1.3 Impact of different sources of errors in calculated variables

From the 100 bootstraps of cuff pressure and MRI measurements, the systematic and random errors in the three calculated hemodynamic variables were calculated (Table 2, *i*=9-11), and 10 of the resulting sampled measurements of the total compliance, total resistance, and maximum elastance of the left ventricle are shown in Figure 5 H-J. Additionally, the sampled uncertainty of the 4D flow MRI-derived variables of the energy loss coefficient of the aortic valve and the compliance of the ascending aorta are shown in Figure 5 F-G.

### 3.2 The confidence intervals from the uncertainty estimation method are validated using a bootstrap approach

We applied the method (Section 2.2.1) on 100 simulated datasets where the truth is known (Section 2.2.3), to evaluate the correctness of the confidence intervals. For the 100 sampled datasets, our PL-based method could find the true biomarker values within the model uncertainty for 91.2±14.6 (mean±sd), median 98%, of the sampled datasets (Figure 6; all found biomarker intervals are presented and compared to the true biomarker value in Supplementary Table S3). The correctness of the confidence interval varied between the different biomarkers, where for example the true value was found 100% of the time for the biomarkers Rpu, Lpv, Rmv, m2LV, and Lav, 94% of the time for Caa and Rao, and only 46 % of the time for the biomarker Emin_LA (Figure 6). The full profiles of all sampled datasets and all investigated biomarkers show well-defined profiles with a mix of identifiable and non-identifiable biomarkers (Supplementary Figure S1). For example, kdiastLA is identifiable because the confidence interval is 23 % of the biomarker value, while Cpvc has a confidence interval that is 2000 % of the biomarker value and thus is unidentifiable. However, to know whether a biomarker is reliable enough for clinical usage, its confidence interval needs to be compared with the cohort variation.

**Figure 6.**
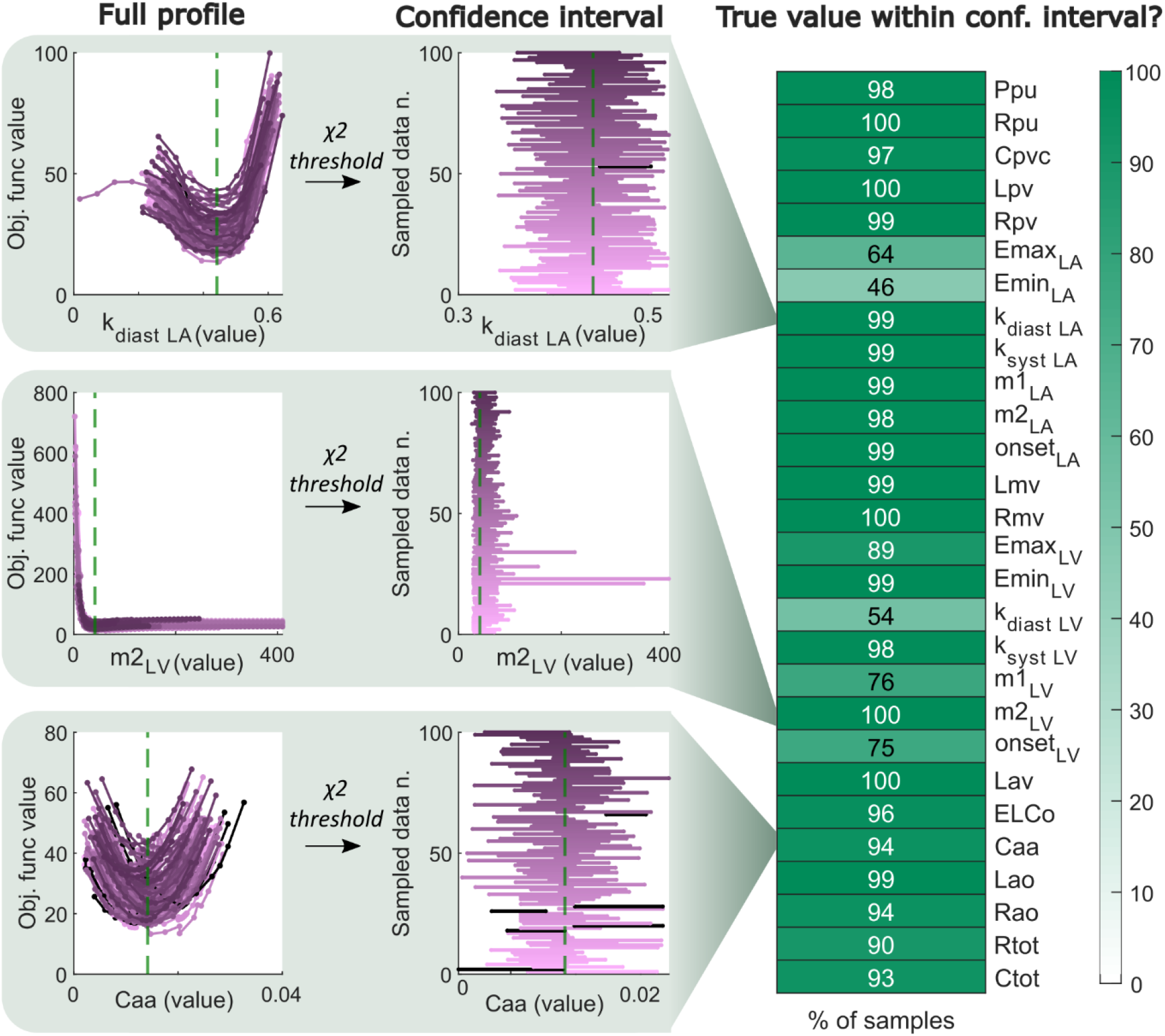
Biomarker uncertainty compared to the true values for all biomarkers. The biomarker uncertainty from profile likelihood for each biomarker and each sampled data resulted in confidence intervals for each biomarker, which is shown for three selected biomarkers to the left. The intervals were defined based on the inverse χ^2^ distribution with one degree of freedom and 95% confidence, and should thus cover the true biomarker value 95% of the time if all assumptions are true. To the right, the % of times that the true biomarker value was within the confidence interval is presented for each biomarker.

### 3.3 Biomarker uncertainty is smaller than the cohort variation for selected biomarkers in a clinical dataset

Finally, the uncertainty estimation method was applied to a clinical dataset from a previous study, and the resulting uncertainties for the six selected model-derived biomarkers are shown in Figure 7. In Figure 7 A-F, the size of the standard deviations of aortic compliance, aortic resistance, left ventricular relaxation rate, maximum elastance of the left atrium, systolic time constant of the left ventricular elastance, and compliance of pulmonary capillaries and veins, are compared between subjects. For all biomarkers except for the compliance of pulmonary capillaries av veins, the uncertainty for the individual subjects is smaller than the cohort variation (Figure 7 G-L). Furthermore, for the aortic compliance, one can see significant differences between almost all of the subjects (Figure 7 A) and a small individual uncertainty (2 %) compared to the cohort variation (36 %) (Figure 7 G), while the individual uncertainty is sorted to be gradually larger compard to the cohort variation for the biomarkers in Figure 7 H-L. Significant differences between subjects along with a relatively small individual uncertainty can also be seen in the biomarkers of aortic resistance, maximum elastance of the left atrium, and left ventricular relaxation rate (Figure 7 B-D, H-J).

**Figure 7.**
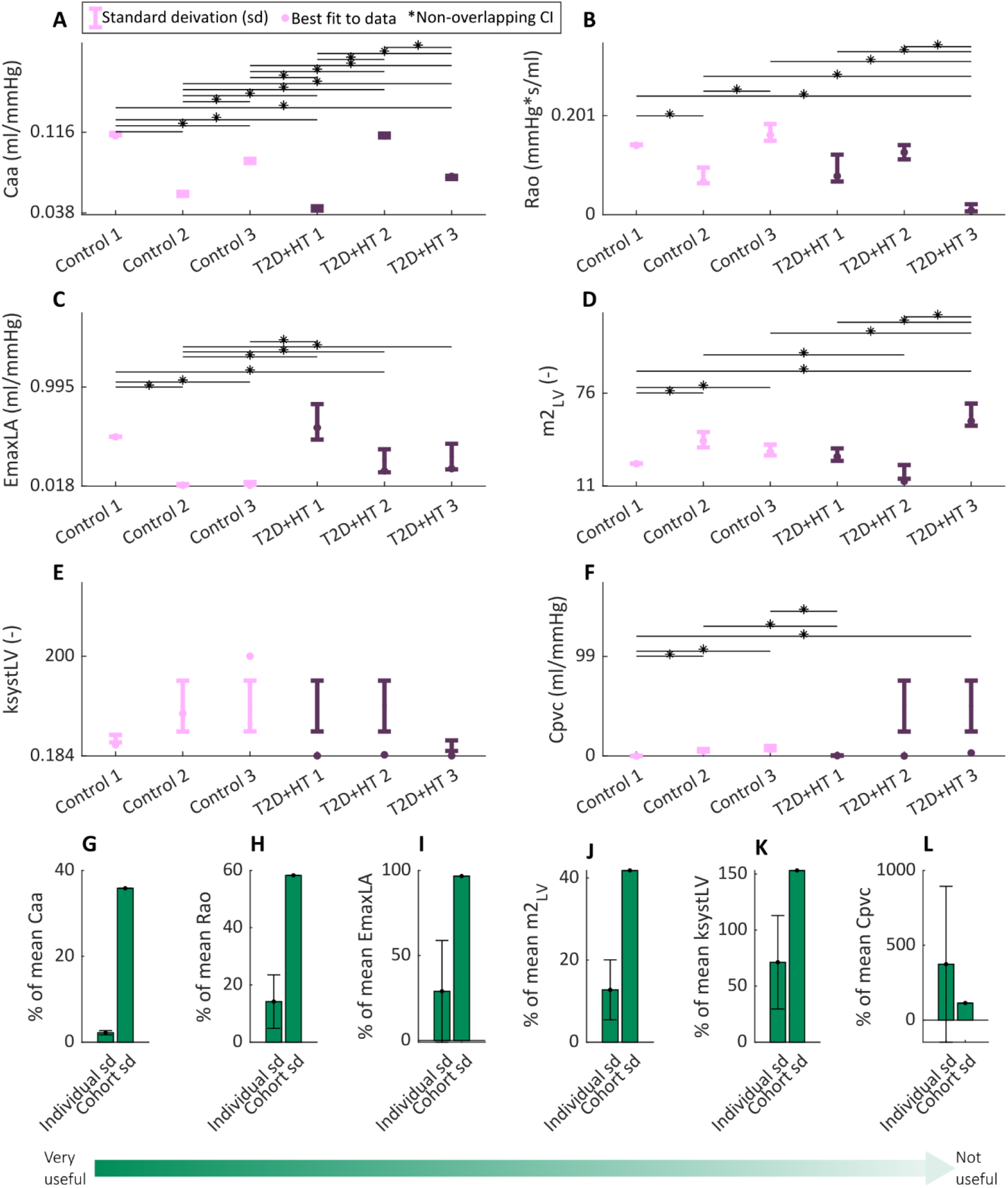
Profile likelihood-derived uncertainty of the patient-specific biomarkers A) Aortic compliance (Caa), B) Aortic resistance (Rao), C) Maximum elastance of the left atrium (Emax LA), D) left ventricular relaxation rate constant (m2LV), E) systolic time constant of the left ventricular elastance function (ksyst LV), and F) compliance of pulmonary capillaries and veins (Cpvc). The standard deviations of the biomarkers are given for six subjects from a clinical cohort (Tunedal et al 2023): three controls (lighter color) and three subjects with both hypertension (HT) and type 2 diabetes (T2D) (darker color), where significant pairwise comparisons are marked with *. In G-L, the individual standard deviation (mean and standard deviation of all subjects) in percent of the mean best biomarker value is compared to the cohort standard deviation in percent of mean best biomarker value for all six biomarkers. The biomarkers are sorted based on their usefulness.

## 4. Discussion

Defining the uncertainty of biomarkers derived from cardiovascular digital twins is crucial for their clinical usefulness. However, the standard approaches to estimating the biomarker uncertainty require a known, additive, and normally distributed data uncertainty, and the errors in hemodynamic data are only partly quantified and do not fulfill these assumptions. To remedy this, we quantified the uncertainty of 4D flow MRI and cuff pressure data (Figure 4, Table 2-3), introduced a novel method to estimate the biomarker uncertainty in a personalized cardiovascular model (Section 2.2.1, Figure 2), and used a bootstrap approach (Figure 3) to show that our method is feasible despite using patient-specific data with many non-normal errors (Figure 6). The method was able to find the true biomarker values within a 95% confidence interval 98% of the time (median), closely following the statistical theoretical value. Thus, our assumptions using the PL method for confidence intervals hold for this simplified combination of normal and non-normal data uncertainty in the cardiovascular model. Additionally, the found uncertainties of e.g. model-derived aortic resistance and left ventricular relaxation rate in a clinical cohort are small enough to be able to compare different groups or compare the same patient in different scenarios – making these biomarkers clinically useful (Figure 7). The presented method can be applied to other data using our modifiable Matlab scripts (https://github.com/kajtu/Uncertainty-estimation, Tunedal 2024).

### 4.1 Estimated data uncertainty

We have herein estimated the uncertainty of patient-specific hemodynamic data based on previous literature. However, some practical simplifications were made, and measurement protocols vary between sites and are continuously being optimized. The true uncertainty for 4D flow MRI, brachial pressure measurements, and the calculated variables might therefore be different from what is estimated here. Further evaluation with new measurements is needed to confirm and assess the effect of the different sources of errors on each measurement.

One example of the estimated uncertainty compared to previous measurements concerns the total flow volume errors in 4D flow MRI. The updated 4D flow MRI consensus statement (Bissell *et al*., 2023) states that all differences in flow volumes within datasets, compared to 2D Flow CMR, and in inter- and intra-observer comparisons ideally should be <= 5% for validation for clinical use, and scan-rescan flow differences should be <10% (Bissell *et al*., 2023). Here, we find that the random errors in stroke volume always are <5%, while the total error including offset and RR smoothing can be as large as 20% in some cases, with a standard deviation of 8.24% in the mitral valve (Figure 4, Table S2). If this is a correct estimation, it would mean that many measurements would be excluded in practice after within and between dataset comparisons if strictly following the consensus statement. However, the estimated total flow volume error of around ±8% is similar to previously reported errors from 4D flow MRI, in the range of ±10% (Jarvis *et al*., 2019; Callaghan *et al*., 2020; Burkhardt *et al*., 2023).

Another example concerns the background offset error. Although the estimated effect of the background offset lacks temporal variation and information on similar errors in blood flow curves close to each other, the effect on the stroke volume is in line with previously reported values. The offset effect on the mass conservation errors in stroke volume of around -0.28-1.51 ± 8.9-11.9 % is similar to the previously found errors in (Chernobelsky *et al*., 2007; Viola *et al*., 2020) of around 6-12% or 7-8 mL. It is, however, a larger offset error than the 0.1±0.5 cm/s or 5% error in cardiac output found in (Hofman *et al*., 2019), but that was an estimation of the background offset alone in 2D phase-contrast MRI, which is typically acquired in the center of the magnet in which the background is smaller, and did not include other sources of errors.

A third example is the RR variability. The RR smoothing error was estimated directly from the flow curves while the real smoothing occurs in k-space and could result in other and possibly smaller effects than seen here. There is a lack of previous studies on the smoothing effect on blood flow throughout the cardiac cycle to confirm the results. Studies comparing to other measurement modalities show that 4D flow MRI often underestimates flow (Callaghan *et al*., 2020; Burkhardt *et al*., 2023), indicating that a smoothing effect exists. Additionally, test-retest studies have found correlations between flow-based parameters and heart rate, speculating that the change in diastolic time might be one of the factors affecting scan-rescan variability (Stoll *et al*., 2018). The diastolic time was indeed found to affect the smoothing error. When the heart rate increases, the time of diastole is shortened while the time of systole is almost constant for heart rates close to rest (Gemignani *et al*., 2008). This results in an underestimation of the early filling peak in the mitral valve when averaging over a range of RR intervals since the early filling of the left ventricle occurs later in the cardiac cycle for higher heart rates (Figure 4A iii). The late filling peak is instead merged with the early filling and increases due to insufficient early filling – resulting in an overestimation when smoothed. The total smoothing effect on the whole cardiac cycle, i.e. the stroke volume, is however close to 0 for all locations.

A fourth example is the estimation of the random errors as purely random. This is also a simplification – for example, the observer variability most likely has a bias where time points close to each other have more similar errors. The estimated random effect results in a spiking in the blood flow curves not often seen in reality (Figure 5), while the random effect on stroke volume around ±1.5% and maximum 5% (Table S2) is in the same magnitude or slightly smaller than previously reported observer variability errors (Markl *et al*., 2011; van Ooij *et al*., 2016; Casas *et al*., 2024).

Finally, the effect of the sources of errors not included in this simplified estimation of measurement uncertainties could be larger than estimated here or lead to other types of uncertainties – such as the effect of respiratory movement on the 4D flow MRI-derived flow curves. It is challenging to isolate specific sources of errors during measurement, and it is therefore also difficult to completely validate the estimated uncertainties. The sampled data uncertainty reported here should be interpreted as that each sampled dataset corresponds to potential values that could result from a specific measurement on a particular patient at a specific time point – not as scan-rescans. Thus, comparisons with other studies are difficult. The final data error is, after all, an estimation done based on previously available measurements.

### 4.2 Evaluation of estimated uncertainty of model-derived biomarkers

Using bootstrapping from the estimated error distributions in data (Figure 3), we showed that our novel method (Figure 2), consisting of: Step 1) estimating combined data uncertainty, Step 2) estimating optimal biomarkers, and Step 3) estimating biomarker uncertainty with PL, can produce reliable confidence intervals from this patient-specific hemodynamic data with non-normal systematic errors (Figure 6). The methods used in Step 2 and Step 3 usually build on the assumption that the data only contains independent, additive, normally distributed noise (Cedersund, 2012). In the case of hemodynamic data, the type of noise that fulfills this assumption is negligible compared to other sources (Table 1). However, if this assumption is fulfilled, this implies that the log-likelihood-based objective function follows a χ^2^ distribution.

In other words, if this assumption is true, the χ^2^ test, used in Step 2, results in rejection or non-rejection of the null hypothesis that the model with the specific parameter vector was used to generate the data. These assumptions also underlie the normal use of PL, used for uncertainty quantification in Step 3. This means that it is not guaranteed that either the χ^2^ test or the profile likelihood method can be used in cases of non-normal data. However, the two methods might still be applicable in specific examples, but this needs to be checked in each example. Our analysis shows that our method, combining all error sources in a combined error ς (Eq 1), applied to 4D flow MRI and cuff pressure data, is an example where the χ^2^ and PL methods are acceptable. More specifically, if our added errors in Step 1 together with our estimated combined data uncertainty still result in the statistics being fulfilled where a 95% confidence interval in practice results in 95% confidence, PL can be used despite the normality assumption. We found that the true value of the biomarker was within the confidence interval in 90-100% of the cases for most biomarkers, showing that the confidence interval was correct and that our method can be used to provide reliable uncertainty estimates for these biomarkers (Figure 6).

The correctness of the confidence interval varied between the biomarkers. For some of the biomarkers, the true value is found within the confidence interval close to 95% of the time, such as Caa and Rao (Figure 6). Some other biomarkers are too well-determined with too small confidence intervals, resulting in a smaller number of found true values, such as 54 % for kdiastLV, 46 % for Emin LA, and 64% for Emax LA. Finally, some of the biomarkers found the true value 100% of the time, indicating a too large uncertainty. These results are comparable to similar analyses when the assumptions in the PL and χ^2^ test are fulfilled, where the agreement between predictions and true states has been reported to be 92-99% (Villaverde *et al*., 2022) and >95% (Kreutz *et al*., 2012) for 95% confidence intervals. Also for other uncertainty estimation methods, such as Markov Chain Monte-Carlo sampling (MCMC), the % of times that the true value lies within the 95% confidence interval can in practice vary between 35-100% for the best-performing algorithms (Valderrama-Bahamóndez & Fröhlich, 2019). In other words, our result validates the usage of the χ^2^ test and PL as described using our method. Note that this also validates the size of the combined errors ς characterized in Step 1.

The approach of combining both the random and systematic errors into ς in Step 1 is a simplification compared to the classical approach of estimating the systematic errors as parameters in the model (Cedersund, 2012). However, the classical approach was implemented for a few biomarkers, where the RR variation was estimated by averaging 5 simulated heartbeats, and in practice the resulting parameter estimation and PL were more computationally heavy than our simplified approach. To reduce the computational time, a third approach, which was not tested, could be to use the empirical function *RR*(*y*_*i*_) instead of simulating the RR error. However, since our approach of combined errors provided reliable confidence intervals for the biomarkers while being faster than the classical approach, this was the method chosen in Step 1.

The PL method was used as the third step in our method of defining biomarker uncertainty. The PL method provides comprehensive confidence intervals independent of unidentifiability, but it is just one out of many other methods to estimate the biomarker uncertainty. For example, SA, MCMC (Raue *et al*., 2013; Schiavazzi *et al*., 2017) parametric bootstrapping (Chang *et al*., 2017; Domogo & Ottesen, 2021) or unscented Kalman filters (Pant *et al*., 2014; Meiburg *et al*., 2020) can be used. However, most of these methods have the same fundamental hypothesis of normally distributed data as the PL method. Among these methods, PL is advantageous since the actual likelihood is sampled based on agreement with the measured data, instead of using pre-determined parameter distributions and without any need for prior identifiability for convergence as for example MCMC does. Additionally, the PL method can be taken further to also estimate the uncertainty of model predictions (Cedersund, 2012; Kreutz *et al*., 2012). This allows for the evaluation of predictions such as of hemodynamics after valve replacement in aortic stenosis patients (Meiburg *et al*., 2020) predictions of the risk of post-hepatectomy portal hypertension (Golse *et al*., 2021) and predictions of cardiac remodeling during different stages of pregnancy (Comunale *et al*., 2021). An example of predictions with uncertainties is given in the Supplementary Figure S2.

### 4.3 Identification of clinically useful model-derived biomarkers

Using the PL approach and the estimated data uncertainty, we could determine the clinical usability of six selected model-derived biomarkers in a clinical cohort. Of these selected biomarkers, the biomarkers m2LV (left ventricular relaxation rate), Caa (aortic compliance), Rao (aortic resistance), and EmaxLA (maximum elastance of the left atrium) had small uncertainty (2-29 %) compared to the cohort variation (36-97 %) (Figure 7 G-J). This allows for comparisons between different subjects within the cohort (Figure 7 A-D). Interestingly, these biomarkers were shown in Tunedal *et al* 2023 to differ on a *group* level between controls and subjects with T2D and/or hypertension, particularly the left ventricular relaxation rate. With these new results, we show that the ventricular relaxation rate is useful on a *patient-specific* level. Additionally, other biomarkers, such as ksyst LV (the systolic time constant of the left ventricular elastance function), cannot be used to differentiate between subjects due to too large uncertainty. In a clinical patient-specific setting, such differentiation between patients, or between different time points for the same patient, is crucial. This quantification of biomarker uncertainty is thus crucial for knowing which model-derived biomarkers that are clinically useful, which previously was unknown due to unknown data uncertainty. However, the usefulness is also highly dependent on the amount of data as well as the uncertainty of the data. Thus, some of the biomarkers with larger uncertainties could become more useful with more or better estimation data. Additionally, only a few selected biomarkers were investigated as proof of concept, and there are most likely several more of the model-derived biomarkers that are well-determined.

#### Limitations

There are a number of limitations of this study. The main limitation of this study is the simplified quantification of data uncertainty. Additionally, only 100 bootstrap samples were used due to the high computational cost of estimating the uncertainty for all 28 biomarkers for each bootstrap. With more samples, the results would be more stabilized (Davidson & MacKinnon, 2000). Furthermore, we cannot be sure that the global minimum was found for all steps in the biomarker profile likelihoods, but the parameter estimation was good enough to see that the PL succeeded in finding robust profiles and reliable confidence intervals. Moreover, the conclusions on biomarker uncertainty found herein are only applicable to the MRI and pressure measurements evaluated here - further evaluation for other modalities and systems is needed if data from other modalities are used to estimate model biomarkers. However, the same method could be applied to new systems, starting with quantifying the patient-specific data uncertainty, and then estimating model uncertainty using our new method. To finally take the next step towards model credibility, the model-derived biomarkers for each application need to be validated with independent data.

## Conclusion

We show that our new method (Figure 2) consisting of Step 1) defining the combined data uncertainty including all non-normal errors, Step 2) finding optimal biomarker values, and Step 3)calculating biomarker uncertainty using PL, together with the quantified non-normal data uncertainty (Table 2-3, Figure 4), provide correct confidence intervals for biomarkers derived from the cardiovascular digital twin (Figure 6), despite simplified methodological assumptions. Additionally, we show that the combined sources of errors in 4D flow MRI and cuff pressure measurements from a clinical cohort result in several biomarker uncertainties that are small enough to compare different subjects and therefore are clinically useful (Figure 7). This method of uncertainty quantification and PL-based uncertainty estimation can be used to provide reliable confidence intervals of model-based biomarkers and predictions, which paves the way for the clinical use of cardiovascular digital twins.

## Acknowledgments

The research is supported by the *Swedish Research Council* (Grant numbers 2018-04454 and 2022-03931, TE; 2018-05418 and 2018-03319, GC), the *Swedish Heart and Lung Foundation* (Grant number 20210441, TE) and the *County Council of Östergötland* (RÖ-987498, TE). GC also acknowledges support from CENIIT (15.09), the Swedish Foundation for Strategic Research (ITM17-0245), SciLifeLab National COVID-19 Research Program financed by the Knut and Alice Wallenberg Foundation (2020.0182), the H2020 project PRECISE4Q (777107), the Swedish Fund for Research without Animal Experiments (F2019-0010), ELLIIT (2020-A12), VINNOVA (VisualSweden, 2020-04711), and the Horizon Europe project STRATIF-AI (101080875). Finally, GC acknowledges scientific support from the Exploring Inflammation in Health and Disease (X‐HiDE) Consortium, which is a strategic research profile at Örebro University funded by the Knowledge Foundation (20200017).

The computations were enabled by resources provided by the National Academic Infrastructure for Supercomputing in Sweden (NAISS) and the Swedish National Infrastructure for Computing (SNIC), partially funded by the Swedish Research Council through grant agreements 2021/3-35.

## Author contributions

K.T, G.C, and T.E participated in the conception and design of the study. K.T performed the literature study and computational analysis and drafted the manuscript. All authors interpreted the results, edited, and revised the manuscript. All authors read and approved the final version of the manuscript.

## Competing interests

The authors declare that they have no competing financial interests.

## Data availability

All code to sample data and use the described method to estimate the biomarker uncertainty is available at https://github.com/kajtu/Uncertainty-estimation (Tunedal, 2024).

## Notes

https://zenodo.org/doi/10.5281/zenodo.13312195

### Competing Interest Statement

The authors have declared no competing interest.

## References

ANSI/AAMI/ISO (2018). Non-invasive sphygmomanometers Part 2: Clinical investigation of intermittent automated measurement type. Available at: https://www.iso.org/standard/73339.html [Accessed May 3, 2023].

Bissell MM et al. (2023). 4D Flow cardiovascular magnetic resonance consensus statement: 2023 update. Journal of Cardiovascular Magnetic Resonance 25, 40.

Bowman AW, Frihauf PA & Kovács SJ (2004). Time-varying effective mitral valve area: prediction and validation using cardiac MRI and Doppler echocardiography in normal subjects. American Journal of Physiology-Heart and Circulatory Physiology 287, H1650–H1657.

Burkhardt BEU, Kellenberger CJ, Callaghan FM, Valsangiacomo Buechel ER & Geiger J (2023). Flow evaluation software for four-dimensional flow MRI: a reliability and validation study. Radiol med; DOI: 10.1007/s11547-023-01697-4.

Callaghan FM, Burkhardt B, Geiger J, Valsangiacomo Buechel ER & Kellenberger CJ (2020). Flow quantification dependency on background phase correction techniques in 4D-flow MRI. Magnetic Resonance in Medicine 83, 2264–2275.

Casas B, Lantz J, Viola F, Cedersund G, Bolger AF, Carlhäll CJ, Karlsson M & Ebbers T (2017). Bridging the gap between measurements and modelling: A cardiovascular functional avatar. Scientific Reports 7, 1–15.

Casas B, Tunedal K, Viola F, Cedersund G, Carlhäll C-J, Karlsson M & Ebbers T (2024). Reproducibility of 4D Flow MRI-based Personalized Cardiovascular Models; Inter-sequence, Intra-observer, and Inter-observer variability. 2024.06.13.597551. Available at: 10.1101/2024.06.13.597551v1 [Accessed June 28, 2024].

Casas B, Viola F, Cedersund G, Bolger AF, Karlsson M, Carlhäll C-J & Ebbers T (2018). Non-invasive Assessment of Systolic and Diastolic Cardiac Function During Rest and Stress Conditions Using an Integrated Image-Modeling Approach. Frontiers in Physiology 9, 1515.

Casciaro ME, Pascaner AF, Guilenea FN, Alcibar J, Gencer U, Soulat G, Mousseaux E & Craiem D (2021). 4D flow MRI: Impact of region of interest size, angulation and spatial resolution on aortic flow assessment. Physiological Measurement 42, 035004.

Cedersund G (2012). Conclusions via unique predictions obtained despite unidentifiability - New definitions and a general method. FEBS Journal 279, 3513–3527.

Chang KC, Dutta S, Mirams GR, Beattie KA, Sheng J, Tran PN, Wu M, Wu WW, Colatsky T, Strauss DG & Li Z (2017). Uncertainty Quantification Reveals the Importance of Data Variability and Experimental Design Considerations for in Silico Proarrhythmia Risk Assessment. Frontiers in Physiology.

Chernobelsky A, Shubayev O, Comeau CR & Wolff SD (2007). Baseline correction of phase contrast images improves quantification of blood flow in the great vessels. Journal of Cardiovascular Magnetic Resonance 9, 681–685.

Chung CS & Kovács SJ (2006). Consequences of Increasing Heart Rate on Deceleration Time, the Velocity–Time Integral, and E/A. The American Journal of Cardiology 97, 130–136.

Comunale G, Susin FM & Mynard JP (2021). Ventricular wall stress and wall shear stress homeostasis predicts cardiac remodeling during pregnancy: A modeling study. International Journal for Numerical Methods in Biomedical Engineeringe3536.

Davidson R & MacKinnon JG (2000). Bootstrap tests: how many bootstraps? Econometric Reviews 19, 55–68.

Dillinger H, Walheim J & Kozerke S (2020). On the limitations of echo planar 4D flow MRI. Magnetic Resonance in Medicine 84, 1806–1816.

Domogo AA & Ottesen JT (2021). Patient-specific parameter estimation: Coupling a heart model and experimental data. Journal of Theoretical Biology 526, 110791.

Dyverfeldt P, Bissell M, Barker AJ, Bolger AF, Carlhäll CJ, Ebbers T, Francios CJ, Frydrychowicz A, Geiger J, Giese D, Hope MD, Kilner PJ, Kozerke S, Myerson S, Neubauer S, Wieben O & Markl M (2015). 4D flow cardiovascular magnetic resonance consensus statement. Journal of Cardiovascular Magnetic Resonance 17, 72.

Dyverfeldt P, Ebbers T & Länne T (2014). Pulse wave velocity with 4D flow MRI: Systematic differences and age-related regional vascular stiffness. Magnetic Resonance Imaging 32, 1266–1271.

Eck VG, Donders WP, Sturdy J, Feinberg J, Delhaas T, Hellevik LR & Huberts W (2016). A guide to uncertainty quantification and sensitivity analysis for cardiovascular applications. International Journal for Numerical Methods in Biomedical Engineering 32, e02755.

Egea JA, Henriques D, Cokelaer T, Villaverde AF, MacNamara A, Danciu DP, Banga JR & Saez-Rodriguez J (2014). MEIGO: An open-source software suite based on metaheuristics for global optimization in systems biology and bioinformatics. BMC Bioinformatics 15, 1–9.

Erbel R (2001). Diagnosis and management of aortic dissection Task Force on Aortic Dissection, European Society of Cardiology. European Heart Journal 22, 1642–1681.

Gemignani V, Bianchini E, Faita F, Giannoni M, Pasanisi E, Picano E & Bombardini T (2008). Assessment of cardiologic systole and diastole duration in exercise stress tests with a transcutaneous accelerometer sensor. In 2008 Computers in Cardiology, pp. 153–156.

Golse N, Joly F, Combari P, Lewin M, Nicolas Q, Audebert C, Samuel D, Allard MA, Sa Cunha A, Castaing D, Cherqui D, Adam R, Vibert E & Vignon-Clementel IE (2021). Predicting the risk of post-hepatectomy portal hypertension using a digital twin: A clinical proof of concept. Journal of Hepatology 74, 661–669.

Gul R & Bernhard S (2015). Parametric uncertainty and global sensitivity analysis in a model of the carotid bifurcation: Identification and ranking of most sensitive model parameters. Mathematical Biosciences 269, 104–116.

Harrod KK, Rogers JL, Feinstein JA, Marsden AL & Schiavazzi DE (2021). Predictive Modeling of Secondary Pulmonary Hypertension in Left Ventricular Diastolic Dysfunction. Frontiers in Physiology; DOI: 10.3389/fphys.2021.666915.

Hofman MBM, Rodenburg MJA, Markenroth Bloch K, Werner B, Westenberg JJM, Valsangiacomo Buechel ER, Nijveldt R, Spruijt OA, Kilner PJ, van Rossum AC & Gatehouse PD (2019). In-vivo validation of interpolation-based phase offset correction in cardiovascular magnetic resonance flow quantification: a multi-vendor, multi-center study. Journal of Cardiovascular Magnetic Resonance 21, 30.

Hofman MBM, Visser FC, Van Rossum AC, Vink GQM, Sprenger M & Westerhof N (1995). In Vivo Validation of Magnetic Resonance Blood Volume Flow Measurements with Limited Spatial Resolution in Small Vessels. Magnetic Resonance in Medicine 33, 778–784.

Hose DR, Lawford PV, Huberts W, Hellevik LR, Omholt SW & van de Vosse FN (2019). Cardiovascular models for personalised medicine: Where now and where next? Medical Engineering and Physics 72, 38–48.

Jarvis K, Schnell S, Barker AJ, Rose M, Robinson JD, Rigsby CK & Markl M (2019). Caval to pulmonary 3D flow distribution in patients with Fontan circulation and impact of potential 4D flow MRI error sources. Magnetic Resonance in Medicine 81, 1205–1218.

Khodaei S, Henstock A, Sadeghi R, Sellers S, Blanke P, Leipsic J, Emadi A & Keshavarz-Motamed Z (2021). Personalized intervention cardiology with transcatheter aortic valve replacement made possible with a non-invasive monitoring and diagnostic framework. Scientific Reports 11, 1–28.

Kim Y-H, Marom EM, Herndon JE & McAdams HP (2005). Pulmonary vein diameter, cross-sectional area, and shape: CT analysis. Radiology 235, 43–49; discussion 49-50.

Kreutz C, Raue A, Kaschek D & Timmer J (2013). Profile likelihood in systems biology. FEBS Journal 280, 2564–2571.

Kreutz C, Raue A & Timmer J (2012). Likelihood based observability analysis and confidence intervals for predictions of dynamic models. BMC Systems Biology; DOI: 10.1186/1752-0509-6-120.

Lewis PS (2019). Oscillometric measurement of blood pressure: a simplified explanation. A technical note on behalf of the British and Irish Hypertension Society. J Hum Hypertens 33, 349–351.

Liu J, Li Y, Li J, Zheng D & Liu C (2022). Sources of automatic office blood pressure measurement error: a systematic review. Physiol Meas 43, 09TR02.

Lövfors W, Simonsson C, Komai AM, Nyman E, Olofsson CS & Cedersund G (2021). A systems biology analysis of adrenergically stimulated adiponectin exocytosis in white adipocytes. Journal of Biological Chemistry 297, 101221.

Macdonald JA, Beshish AG, Corrado PA, Barton GP, Goss KN, Eldridge MW, François CJ & Wieben O (2020). Feasibility of Cardiovascular Four-dimensional Flow MRI during Exercise in Healthy Participants. Radiology: Cardiothoracic Imaging 2, e190033.

Markl M, Frydrychowicz A, Kozerke S, Hope M & Wieben O (2012). 4D flow MRI. Journal of Magnetic Resonance Imaging 36, 1015–1036.

Markl M, Wallis W & Harloff A (2011). Reproducibility of flow and wall shear stress analysis using flow-sensitive four-dimensional MRI. Journal of Magnetic Resonance Imaging 33, 988–994.

Meiburg R, Huberts W, Rutten MCM & van de Vosse FN (2020). Uncertainty in model-based treatment decision support: Applied to aortic valve stenosis. International Journal for Numerical Methods in Biomedical Engineering 36, e3388.

Montalba C, Urbina J, Sotelo J, Andia ME, Tejos C, Irarrazaval P, Hurtado DE, Valverde I & Uribe S (2018). Variability of 4D flow parameters when subjected to changes in MRI acquisition parameters using a realistic thoracic aortic phantom. Magnetic Resonance in Medicine 79, 1882–1892.

Musuamba FT et al. (2021). Scientific and regulatory evaluation of mechanistic in silico drug and disease models in drug development: Building model credibility. CPT: Pharmacometrics and Systems Pharmacology 10, 804–825.

van Ooij P, Powell AL, Potters WV, Carr JC, Markl M, & Barker, and Alex J. (2016). Reproducibility and interobserver variability of systolic blood flow velocity and 3D wall shear stress derived from 4D flow MRI in the healthy aorta. Journal of Magnetic Resonance Imaging 43, 236–248.

Pant S, Fabrèges B, Gerbeau J-F & Vignon-Clementel IE (2014). A methodological paradigm for patient-specific multi-scale CFD simulations: from clinical measurements to parameter estimates for individual analysis. Int J Numer Method Biomed Eng 30, 1614–1648.

Pathmanathan P, Cordeiro JM & Gray RA (2019). Comprehensive Uncertainty Quantification and Sensitivity Analysis for Cardiac Action Potential Models. Frontiers in Physiology 10, 721.

Pravdivtseva MS, Gaidzik F, Berg P, Ulloa P, Larsen N, Jansen O, Hövener J-B & Salehi Ravesh M (2022). Influence of Spatial Resolution and Compressed SENSE Acceleration Factor on Flow Quantification with 4D Flow MRI at 3 Tesla. Tomography 8, 457–478.

Quicken S, Donders WP, van Disseldorp EMJ, Gashi K, Mees BME, van de Vosse FN, Lopata RGP, Delhaas T & Huberts W (2016). Application of an Adaptive Polynomial Chaos Expansion on Computationally Expensive Three-Dimensional Cardiovascular Models for Uncertainty Quantification and Sensitivity Analysis. Journal of Biomechanical Engineering; DOI: 10.1115/1.4034709.

Randall EB, Randolph NZ, Alexanderian A & Olufsen MS (2021). Global sensitivity analysis informed model reduction and selection applied to a Valsalva maneuver model. Journal of Theoretical Biology 526, 110759.

Raue A, Kreutz C, Theis FJ & Timmer J (2013). Joining forces of Bayesian and frequentist methodology: A study for inference in the presence of non-identifiability. Philosophical Transactions of the Royal Society A: Mathematical, Physical and Engineering Sciences; DOI: 10.1098/rsta.2011.0544.

Roes SD, Korosoglou G, Schär M, Westenberg JJ, van Osch MJP, de Roos A & Stuber M (2008). Correction for heart rate variability during 3D whole heart MR coronary angiography. Journal of Magnetic Resonance Imaging 27, 1046–1053.

de Roquefeuil M, Vuissoz P-A, Escanyé J-M & Felblinger J (2013). Effect of physiological Heart Rate variability on quantitative T2 measurement with ECG-gated Fast Spin Echo (FSE) sequence and its retrospective correction. Magnetic Resonance Imaging 31, 1559–1566.

Santini F, Pansini M, Hrabak-Paar M, Yates D, Langenickel TH, Bremerich J, Bieri O & Schubert T (2020). On the optimal temporal resolution for phase contrast cardiovascular magnetic resonance imaging: establishment of baseline values. Journal of Cardiovascular Magnetic Resonance 22, 72.

Schiavazzi DE, Baretta A, Pennati G, Hsia TY & Marsden AL (2017). Patient-specific parameter estimation in single-ventricle lumped circulation models under uncertainty. International Journal for Numerical Methods in Biomedical Engineering; DOI: 10.1002/cnm.2799.

Stergiou GS, Palatini P, Parati G, O’Brien E, Januszewicz A, Lurbe E, Persu A, Mancia G & Kreutz R (2021). 2021 European Society of Hypertension practice guidelines for office and out-of-office blood pressure measurement. Journal of Hypertension 39, 1293.

Stoll VM, Loudon M, Eriksson J, Bissell MM, Dyverfeldt P, Ebbers T, Myerson SG, Neubauer S, Carlhäll CJ & Hess AT (2018). Test-retest variability of left ventricular 4D flow cardiovascular magnetic resonance measurements in healthy subjects. Journal of Cardiovascular Magnetic Resonance 20, 1–10.

Tunedal K (2024). Uncertainty code. Available at: 10.5281/zenodo.13312195.

Tunedal K, Viola F, Garcia BC, Bolger A, Nyström FH, Östgren CJ, Engvall J, Lundberg P, Dyverfeldt P, Carlhäll C-J, Cedersund G & Ebbers T (2023). Haemodynamic effects of hypertension and type 2 diabetes: Insights from a 4D flow MRI-based personalized cardiovascular mathematical model. The Journal of Physiology 601, 3765–3787.

Valderrama-Bahamóndez GI & Fröhlich H (2019). MCMC Techniques for Parameter Estimation of ODE Based Models in Systems Biology. Front Appl Math Stat; DOI: 10.3389/fams.2019.00055.

Viceconti M, Pappalardo F, Rodriguez B, Horner M, Bischoff J & Musuamba Tshinanu F (2021). In silico trials: Verification, validation and uncertainty quantification of predictive models used in the regulatory evaluation of biomedical products. Methods 185, 120–127.

Villaverde AF, Raimúndez E, Hasenauer J & Banga JR (2022). Assessment of Prediction Uncertainty Quantification Methods in Systems Biology. IEEE/ACM Transactions on Computational Biology and Bioinformatics1–12.

Viola F, Dyverfeldt P, Carlhäll CJ & Ebbers T (2020). Data Quality and Optimal Background Correction Order of Respiratory-Gated k-Space Segmented Spoiled Gradient Echo (SGRE) and Echo Planar Imaging (EPI)-Based 4D Flow MRI. Journal of Magnetic Resonance Imaging 51, 885– 896.

Wan Y, Heneghan C, Stevens R, McManus RJ, Ward A, Perera R, Thompson M, Tarassenko L & Mant D (2010). Determining which automatic digital blood pressure device performs adequately: a systematic review. J Hum Hypertens 24, 431–438.

Wentland AL, Grist TM & Wieben O (2014). Review of MRI-based measurements of pulse wave velocity: a biomarker of arterial stiffness. Cardiovascular Diagnosis and Therapy 4, 193–206.

Wittkampf FHM, Vonken E-J, Derksen R, Loh P, Velthuis B, Wever EFD, Boersma LVA, Rensing BJ & Cramer M-J (2003). Pulmonary Vein Ostium Geometry. Circulation 107, 21–23.

Wolf RL, Ehman RL, Riederer SJ & Rossman PJ (1993). Analysis of systematic and random error in MR volumetric flow measurements. Magnetic Resonance in Medicine 30, 82–91.

Yamamoto K, Masuyama T, Tanouchi J, Doi Y, Kondo H, Hori M, Kitabatake A & Kamada T (1993). Effects of heart rate on left ventricular filling dynamics: assessment from simultaneous recordings of pulsed Doppler transmitral flow velocity pattern and haemodynamic variables. Cardiovascular Research 27, 935–941.

