## Supplementary for "Uncertainty in cardiovascular digital twins despite non-normal errors in 4D flow MRI: identifying reliable biomarkers such as ventricular relaxation rate"

This supplementary information contains additional methods, tables, and figures that might be of interest to the reader.

**Method**

*Table S1. Description and units of all estimated model parameters in the model from Tunedal et al 2023 and Casas et al 2018.*

| **Parameter name** | **Unit** | **Description** |
| --- | --- | --- |
| Caa* | mL/mmHg | Capacitance of the aorta |
| Cpvc | mL/mmHg | Capacitance of pulmonary capillaries and veins |
| Ctot* | mL/mmHg | Total compliance of the system |
| ELCo* | cm^2 | Energy loss coefficient of the aortic valve |
| Emax_LA | mmHg/mL | Maximal elastance of the LA |
| Emax_LV* | mmHg/mL | Maximal elastance of the LV |
| Emin_LA | mmHg/mL | Minimal (passive) elastance of the LA |
| Emin_LV | mmHg/mL | Minimal (passive) elastance of the LV |
| k_diast_LA | s | Diastolic time constant of the LA |
| k_diast_LV | s | Diastolic time constant of the LV |
| k_syst_LA | s | Systolic time constant of the LA |
| k_syst_LV | s | Systolic time constant of the LV |
| Lao | mmHg*s^2/mL | Inertance of the ascending aorta |
| Lav | mmHg*s^2/mL | Inertance of the aortic valve |
| Lmv | mmHg*s^2/mL | Inertance of the mitral valve |
| Lpv | mmHg*s^2/mL | Inertance of the pulmonary veins |
| m1_LA | - | Contraction rate constant of the LA |
| m1_LV | - | Contraction rate constant of the LV |
| m2_LA | - | Relaxation rate constant of the LA |
| m2_LV | - | Relaxation rate constant of the LV |
| onset_LA | fraction of T | Onset of contraction of the LA |
| onset_LV | fraction of T | Onset of contraction of the LV |
| Ppu | mmHg | Pulmonary capillary pressure |
| Rao | mmHg·s/mL | Resistance of the ascending aorta |
| Rmv | mmHg·s/mL | Resistance of the mitral valve |
| Rpu | mmHg·s/mL | Resistance of pulmonary capillaries |
| Rpv | mmHg·s/mL | Resistance of pulmonary veins |
| Rtot* | mmHg·s/mL | Total resistance of the system |

**Results**

*Table S2. Quantification of the errors in stroke volume in the mitral valve (MV), aortic valve (AV), ascending aorta (AA), and pulmonary veins (PV) compared to the true value, and stroke volume differences between the different locations. Mean±standard deviations are reported for all errors.*

|  | Total error (mL) | Total error (%) | Random errors only (%) | Systematic errors only (%) |
| --- | --- | --- | --- | --- |
| MV | 0.37 ± 4.87 | 0.63 ± 8.24 | 0.19 ± 1.51 | 0.43 ± 8.19 |
| AV | 0.11 ± 3.23 | 0.18 ± 5.48 | -0.17 ± 1.60 | 0.36 ± 5.70 |
| AA | -0.62 ± 4.89 | -1.05 ± 8.28 | -0.12 ± 1.55 | -0.93 ± 8.15 |
| PV | -0.18 ± 3.96 | -0.31 ± 6.74 | -0.02 ± 1.44 | -0.29 ± 6.52 |
| MV-AV | 0.41 ± 5.43 | 0.50 ± 9.20 | 0.61 ± 2.22 | 0.15 ± 9.34 |
| MV-AA | 1.07 ± 7.10 | 1.83 ± 12.11 | 0.45 ± 2.23 | 1.51 ± 11.92 |
| MV-PV | 0.92 ± 5.82 | 1.46 ± 9.95 | 0.84 ± 2.07 | 1.23 ± 9.51 |
| AV-AA | 0.67 ± 6.01 | 1.34 ± 10.43 | -0.16 ± 2.37 | 1.37 ± 10.15 |
| AV-PV | 0.52 ± 5.22 | 0.96 ± 8.99 | 0.23 ± 2.07 | 1.09 ± 8.91 |
| AA-PV | -0.15 ± 6.24 | -0.38 ± 10.88 | 0.39 ± 1.90 | -0.28 ± 10.54 |

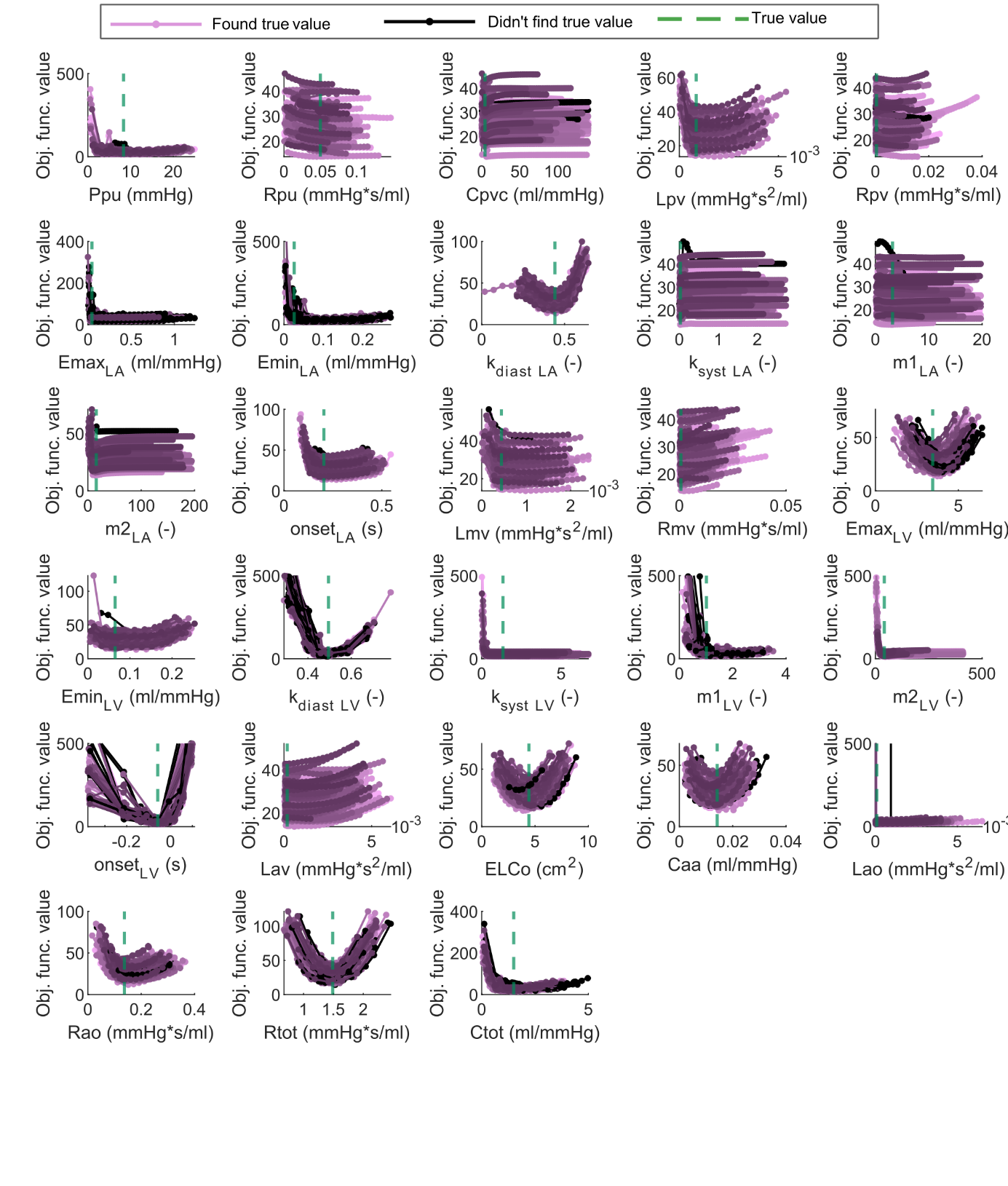
 *Figure S1. Biomarker uncertainty profiles for all 100 sampled datasets and all 28 estimated biomarkers in the model. Purple curves indicate that the true value (green dashed line) was found within the model uncertainty, while black curves indicate that the true value was not found.*

*Table S3. Estimated biomarker uncertainty compared to the true value for all estimated biomarkers in the model. For each biomarker, the minimum and mean ± standard deviation of all lower bounds and the maximum and mean ± standard deviation of the upper bounds of all samples are given. The percentage of identifiability is given in the right columns, where yes corresponds to identifiable, semi corresponds to semi-identifiable (where either the lower or the upper bound was found), and no corresponds to non-identifiability.*

|  | Lower value:  mean ± sd (min) | Upper value:  mean ± sd (max) | True value | Identifiable? | | | |
| --- | --- | --- | --- | --- | --- | --- | --- |
|  |  |  |  | Yes (%) | Semi: only lower bound found (%) | Semi: only upper bound found (%) | No (%) |
| Ppu | 0.042 *±* 0.010 (0.020) | 0.155 *±* 0.018 (0.182) | 0.065 | 78 | 2 | 13 | 7 |
| Rpu | 0.014 ± 0.010 (0.000) | 0.079 ± 0.019 (0.147) | 0.050 | 0 | 33 | 0 | 67 |
| Cpvc | 4.111 ± 1.436 (1.944) | 15.027 ± 2.361 (18.953) | 8.332 | 16 | 13 | 6 | 65 |
| Lpv | 0.001 ± 0.000 (0.000) | 0.003 ± 0.000 (0.004) | 0.001 | 57 | 26 | 10 | 7 |
| Rpv | 1.826 ± 1.268 (0.162) | 90.728 ± 43.95 (139.95) | 4.442 | 0 | 1 | 2 | 97 |
| Emax_LA | 0.017 ± 0.007 (0.002) | 1.696 ± 0.791 (2.74) | 0.027 | 45 | 13 | 38 | 4 |
| Emin_LA | 8.096 ± 1.233 (4.422) | 91.5 ± 57.1 (194.4) | 15.67 | 79 | 9 | 10 | 2 |
| k_diast_LA | 0.968 ± 0.123 (0.574) | 1.654 ± 0.312 (2.438) | 1.015 | 96 | 1 | 3 | 0 |
| k_syst_LA | -0.071 ± 0.010 (-0.090) | -0.043 ± 0.014 (-0.007) | -0.058 | 0 | 0 | 5 | 95 |
| m1_LA | 3.755 ± 0.397 (2.582) | 5.234 ± 0.512 (6.407) | 4.431 | 0 | 1 | 0 | 99 |
| m2_LA | 0.000 ± 0.000 (0.000) | 0.002 ± 0.001 (0.005) | 0.000 | 39 | 51 | 1 | 9 |
| onset_LA | 1.370 ± 0.081 (1.166) | 1.552 ± 0.065 (1.902) | 1.489 | 78 | 14 | 0 | 8 |
| Lmv | 3.178 ± 0.322 (2.401) | 4.142 ± 0.366 (5.190) | 3.451 | 1 | 68 | 1 | 30 |
| Rmv | 0.375 ± 0.223 (0.215) | 4.173 ± 1.374 (6.716) | 1.345 | 0 | 0 | 28 | 72 |
| Emax_LV | 0.526 ± 0.822 (0.119) | 12.894 ± 4.67 (19.75) | 3.214 | 92 | 0 | 8 | 0 |
| Emin_LV | 0.174 ± 0.011 (0.151) | 0.355 ± 0.044 (0.456) | 0.204 | 86 | 5 | 4 | 5 |
| k_diast_LV | 32.62 ± 3.518 (26.15) | 82.0 ± 48.7 (409.5) | 41.55 | 100 | 0 | 0 | 0 |
| k_syst_LV | 0.000 ± 0.000 (0.000) | 0.003 ± 0.001 (0.005) | 0.000 | 0 | 86 | 0 | 14 |
| m1_LV | 0.011 ± 0.002 (0.006) | 0.017 ± 0.002 (0.022) | 0.014 | 95 | 4 | 1 | 0 |
| m2_LV | 0.118 ± 0.012 (0.087) | 0.203 ± 0.030 (0.285) | 0.137 | 93 | 1 | 1 | 5 |
| onset_LV | 1.162 ± 0.330 (0.563) | 1.978 ± 0.382 (3.439) | 1.502 | 100 | 0 | 0 | 0 |
| Lav | 0.001 ± 0.000 (0.000) | 0.017 ± 0.006 (0.034) | 0.001 | 0 | 0 | 55 | 45 |
| ELCo | 0.034 ± 0.015 (0.015) | 0.141 ± 0.032 (0.198) | 0.026 | 89 | 3 | 8 | 0 |
| Caa | 0.386 ± 0.025 (0.340) | 0.490 ± 0.021 (0.522) | 0.442 | 93 | 0 | 4 | 3 |
| Lao | 0.000 ± 0.000 (0.000) | 0.002 ± 0.000 (0.002) | 0.000 | 16 | 3 | 72 | 9 |
| Rao | 0.468 ± 0.012 (0.437) | 0.496 ± 0.007 (0.515) | 0.495 | 94 | 2 | 1 | 3 |
| Rtot | 0.000 ± 0.000 (0.000) | 0.014 ± 0.006 (0.031) | 0.000 | 99 | 0 | 1 | 0 |
| Ctot | 0.055 ± 0.037 (0.020) | 0.464 ± 0.193 (0.876) | 0.044 | 93 | 5 | 2 | 0 |

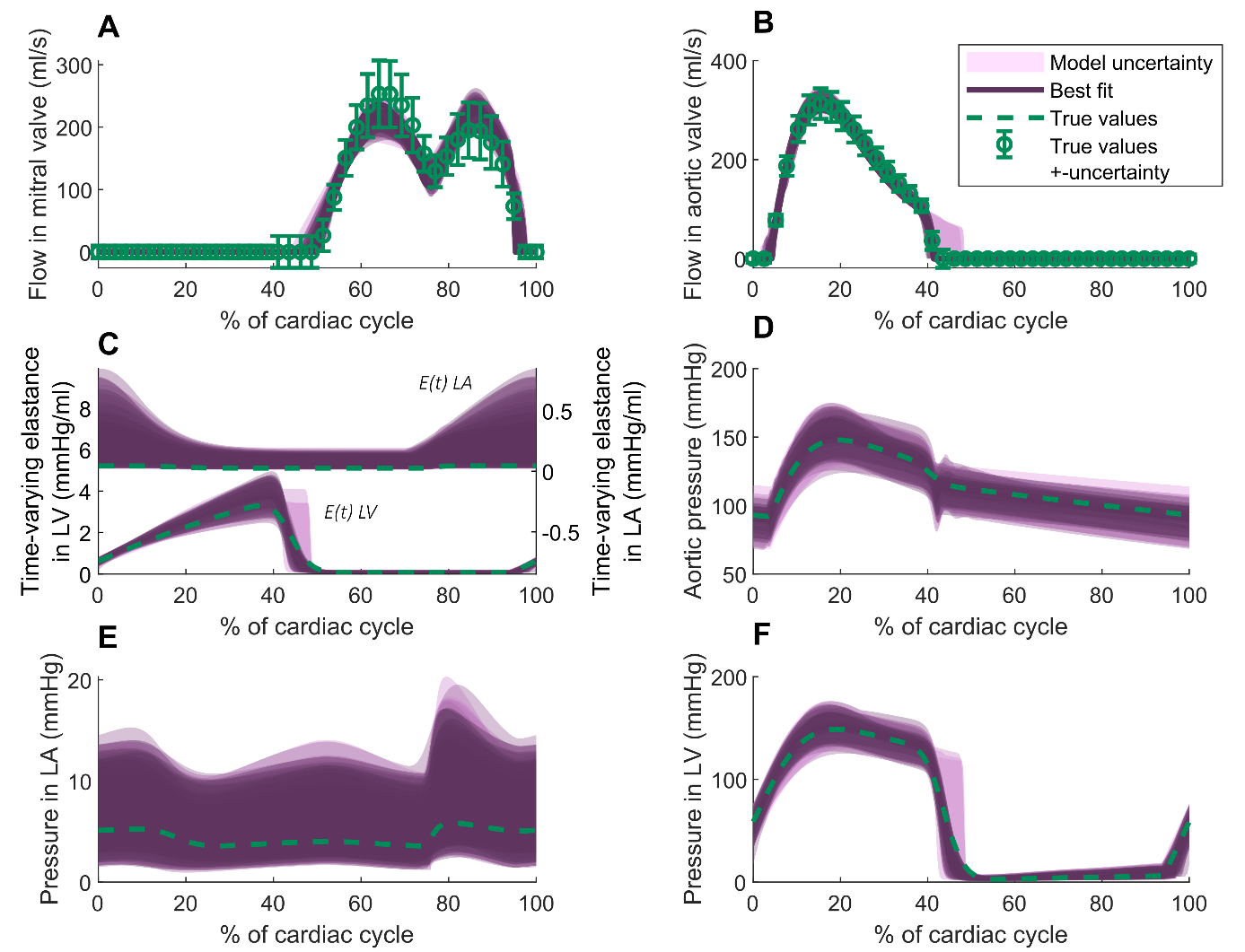
 *Figure S2. Model-derived predictions of A) Blood flow in the mitral valve, B) blood flow in the aortic valve, C) time-varying elastance in the left atrium (LA) and left ventricle (LV), pressure in the D) aorta, E) left atrium and F) left ventricle. An estimate of the uncertainty of each prediction is shown in purple areas, where the uncertainty was derived by simulating all parameter sets collected during the profile likelihood for the biomarkers. The true simulation is shown as a green dashed line, and the true simulation with estimated combined uncertainty is shown in error bars in A and B.*
